# Molecular clock dating under mixture substitution models indicates contemporaneous emergence of *Bradyrhizobium* nodulation and legumes anchored by non-symbiotic relatives

**DOI:** 10.64898/2026.05.28.728399

**Authors:** Sishuo Wang, Lu Ling, Haiwei Luo

**Affiliations:** Simon F.S. Li Marine Science Laboratory, School of Life Sciences and State Key Laboratory of Agrobiotechnology, The Chinese University of Hong Kong, Shatin, Hong Kong SAR; Department of Microbiology, Faculty of Medicine, The Chinese University of Hong Kong, Hong Kong SAR; Institute of Environment, Energy and Sustainability, The Chinese University of Hong Kong, Shatin, Hong Kong SAR

**Keywords:** *Bradyrhizobium*, molecular clock, divergence time, mutualism evolution, mixture model

## Abstract

The transition from free-living bacteria to mutualistic symbionts is a major evolutionary innovation, but its timing and stepwise nature remain unresolved because non-symbiotic ancestors have rarely been sampled. Here we isolated 10 *Bradyrhizobium* strains from non-legume plants and soils, and discovered three basal clades within the *B. japonicum* supergroup that completely lack nodulation (*nod*) genes, which are required for Nod factor synthesis and legume symbiosis. To date the emergence of symbiosis, we employed a mitochondrial endosymbiosis-based molecular clock dating framework that jointly estimates the divergence times of bacterial symbionts and their hosts in a single analysis. Using these non-nodulating lineages as evolutionary anchors, we show that nodulating *B. japonicum* arose contemporaneously with nodulating legumes most likely around 90-80 million years ago. Excluding these basal clades makes rhizobial nodulation appear to predate nodulating legumes, while the use of the poorly-fitting homogeneous models produces a similar temporal distortion. The inferred patterns are robust to alternative calibration strategies, gene sets, and mixture models. Together, these findings highlight the importance of comprehensive taxon sampling and appropriate substitution models in relaxed molecular clock dating, and support a co-emergence model in the origin of legume-rhizobia mutualism.

## Introduction

The symbiosis between *Bradyrhizobium* bacteria and legume plants, defined as the mutualistic relationship in which bacteria induce root nodules where they fix atmospheric nitrogen, underscores the evolutionary significance of nodulation (the process of forming nodules). This symbiosis contributes to agricultural sustainability by enhancing nitrogen fixation, reducing reliance on chemical fertilizers, and promoting soil health (Kebede 2021). Unlike other rhizobia groups, members of *Bradyrhizobium* display highly diverse lifestyles and fall into seven phylogenetic supergroups (phylogroups) (Avontuur et al. 2019; Ormeño-Orrillo and Martínez-Romero 2019). Here, “supergroup” refers to a deeply branching lineage within the genus that is genetically and ecologically distinct. This diversity includes a supergroup formerly known as the photosynthetic supergroup and now called the “*nod*-free but nodulating” supergroup (Ling et al. 2025), which uses a *nod*-independent pathway to nodulate legumes of the genus *Aeschynomene* (Giraud et al. 2007; Okazaki et al. 2016). Here, *nod* genes refer to the bacterial genes that encode the molecular signals (Nod factors) required for nodulation. Other aspects of *Bradyrhizobium* diversity include the widespread distribution of non-symbiotic members in soils (Delgado-Baquerizo et al. 2018; Tao et al. 2021), associations with non-leguminous plants (Wasai-Hara et al. 2020), and loss of symbiotic function during evolution (Sachs et al. 2011). Collectively, these observations point to complex interactions that extend beyond conventional rhizobia-legume symbiosis.

Among non-legumes, only the tropical woody genus *Parasponia* (Cannabaceae, Rosales) forms nitrogen-fixing nodules with *Bradyrhizobium* using a pathway similar to that of legumes (Doyle 2016; van Velzen et al. 2018). The existence of *Parasponia* raises a question: did nodulation capacity originate once in *Bradyrhizobium* before the divergence of legumes and *Parasponia*, or did it arise later and independently in both lineages? Answering this question requires a precise evolutionary timeline that can place the origin of nodulation in *Bradyrhizobium* relative to the diversification of related plant lineages.

*Bradyrhizobium* is presumably the most ancient and genetically diversified rhizobial lineage (Provorov and Andronov 2016; Sprent et al. 2017; Ormeño-Orrillo and Martínez-Romero 2019), making it a promising system for timing the origin of nodulation. A previous study attempted to date this origin (Wang et al. 2020), but its resolution was limited for several reasons. Wang et al. 2020 included only seven *Bradyrhizobium* strains, and similarly few from each other rhizobial genera. It relied on very distantly related cyanobacterial fossils in molecular dating, did not jointly estimate the origin times of rhizobia and nodulating legumes (Caesalpinioideae and Papilionoideae, Fabales) using the same dataset, and lacked non-nodulating basal lineages that could constrain the timing of nodulation from within the *Bradyrhizobium* phylogeny. Further, like most molecular dating studies of that period, Wang et al. 2020 applied only homogeneous substitution models, but recent advances in model development, including our own works, highlight the importance of incorporating mixture models in molecular dating (Demotte et al. 2025; Wang and Luo 2025; Wang and Meade 2026). Consequently, the divergence time estimated in previous studies may need to be revisited with more complex and biologically realistic models.

Most previous studies on *Bradyrhizobium* (and other rhizobia) focused on agriculturally important strains isolated from legumes. Such biased taxon samplings can distort phylogenetic inference, potentially obscuring patterns of evolutionary divergence and misinterpretations of timing. Broader sampling of soil *Bradyrhizobium* revealed more frequent transitions between nodulating and non-nodulating lifestyles than previously recognized (Tao et al. 2021; Bueno de Mesquita et al. 2025; Ling et al. 2026). Recent work also showed that *nod*-independent nodulation with some tropical legumes the *Aeschynomene* genus is widespread in *Bradyrhizobium* from rice paddies (Ling et al. 2025). Together, these findings indicate that sampling non-legume-associated lineages is essential for reconstructing the true evolutionary history of nodulation. An unresolved question is whether successful nodulation evolved through reciprocal coevolution with legumes.

Here we isolated diverse *Bradyrhizobium* strains from non-legume plants and soils. We identified three non-nodulating clades that lack *nod* genes basal to other known members of the *B. japonicum* supergroup, which contains the vast majority of known legume-nodulating strains within the genus (Avontuur et al. 2019; Ormeño-Orrillo and Martínez-Romero 2019). Using these lineages as evolutionary anchors in relaxed molecular clock analyses, we found that nodulating *B. japonicum* supergroup lineages arose contemporaneously with nodulating legumes. The relative timing of these two events was stable across alternative fossil calibrations only when these basal non-symbiotic clades were included; excluding them falsely made rhizobial nodulation appear older than the legumes. We further extended a method for molecular dating we developed in a previous study (Wang and Luo 2025) to support more mixture substitution models, and systematically evaluated their effects on divergence time estimates of the origin of nodulating *Bradyrhizobium* and legumes.

## Results

### Three basal non-nodulating clades in the *B. japonicum* supergroup isolated from diverse habitats

We sequenced the genomes of 10 *Bradyrhizobium* isolates obtained from three non-legume plants: rice, forest trees, and grassland species (Data S1-S2; Fig. S1). Phylogenomic analysis with public *Bradyrhizobium* genomes using 123 orthologous genes shared across the genus identified in our previous study (Tao et al. 2021) consistently placed newly isolated strains, together with 13 strains reported in our recent work (Ling et al. 2026) and three by others, as lineages basal to the known *B. japonicum* supergroup, under the site-compositionally heterogeneous model LG+C20+G with mixture weights estimated using the “-mwopt” option (Fig. 1a). These strains formed three clades, with the earliest-splitting clade containing 4 strains all of which were isolated from the present study, and the other two consisting of 12 and 10 members, respectively (Figs. 1a, S1). This phylogenetic position was consistently recovered under alternative homogeneous substitution models using the full genome set, and under more complex heterogeneous models with a subsampling of genomes (Fig. S2). To improve phylogenetic resolution, we identified 891 single-copy genes conserved across the *B. japonicum* supergroup and repeated the phylogenomic analysis under various substitution models, which confirmed the above pattern (Fig. S3).

**Figure 1.**
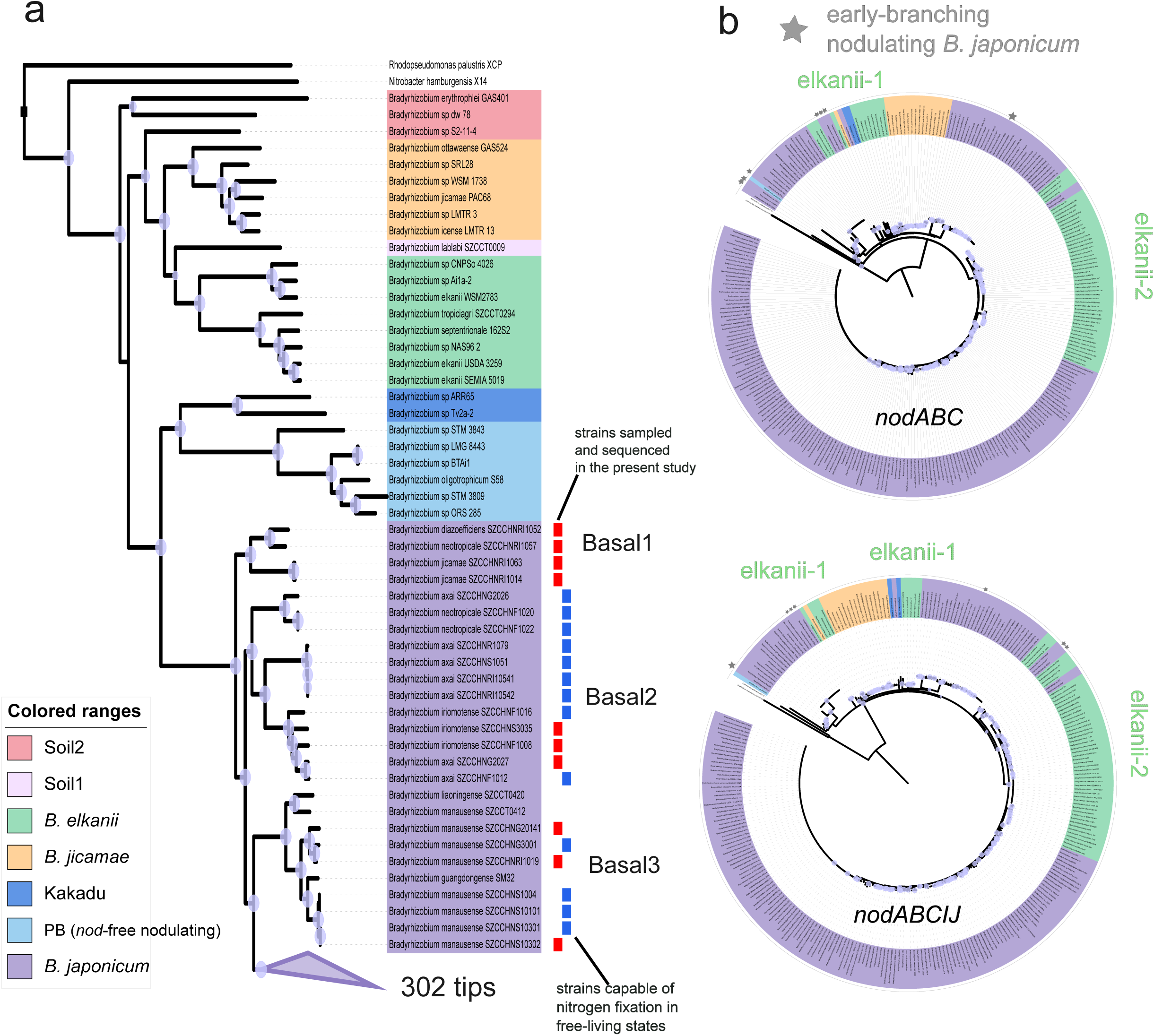
Phylogenetic analysis of *Bradyrhizobium japonicum* supergroup with newly isolated non-nodulating lineages. a) Phylogenomic reconstruction of the *B. japonicum* supergroup based on 123 single-copy genes conserved in the genus constructed under the substitution model LG+C20+G with a posterior mean site frequency (PMSF) approximation (for computational efficiency, LG+C60+G is applied to a subsampling of genomes [Fig. S2]). Strains isolated in the present study are indicated by red squares adjacent to the name of the strain. Blue squares indicate those that have been shown to be capable of nitrogen fixation in a free-living state by a parallel study (Ling et al. 2026) while others are not tested. b) The gene phylogeny of *nodABC* and *nodABCIJ* within *Bradyrhizobium* constructed under LG+C60+G with PMSF. Early-branching nodulating *B. japonicum* strains (Ec3.3, BR10245, BR10247, Cp5.3, AS23.2, NAS80.1, aSej3) are indicated by a grey star. The tree is rooted by outgroup sequences from *Azorhizobium* (rooting with an outgroup-free method points to the same root position). Circles on the nodes denote ultrafast bootstrap values of 90-100, with circle size increasing from smallest to largest.

None of these basal lineages carried *nod* genes. To date, *nod-*independent nodulation has been found in only the “*nod*-free but nodulating” supergroup (Giraud et al. 2007; Ling et al. 2025). Therefore, all newly isolated *B. japonicum* strains are almost certainly non-nodulating, free-living lineages that represent extant relatives of the ancestors of nodulating *B. japonicum*. A parallel study (Ling et al. 2026) independently demonstrates that several strains from this collection (Fig. 1a, tips labeled by a blue square), together with other free-living *Bradyrhizobium* isolates, carry a conserved *nif* island architecture including the oxygen-protective gene *glbO*, supporting their capacity for nitrogen fixation in the free-living state. A separate phylogenetic analysis of *nod* genes (Fig. 1b) identified four origins of nodulation in *Bradyrhizobium*: one within the *B. japonicum* supergroup, one within the *B. jicamae* supergroup, and two within the *B. elkanii* supergroup (we did not analyze the Kakadu supergroup due to very few strains available), broadly consistent with ancestral state reconstruction analysis (Fig. S4). Early-diverging lineages of nodulating *B. japonicum* strains (labeled by grey stars in Fig. 1b) occupied the basal positions in the *nod* gene tree (Fig. 1b). Nodulating strains from *B. jicamae* were nested within *B. japonicum* members, followed by two origins within *B. elkanni*. This pattern suggests HGT of *nod* genes from early-diverging nodulating *B. japonicum* to other *Bradyrhizobium* supergroups, consistent with a previous report (Teulet et al. 2020). The tree topology was robust to substitution model choice (Fig. S5).

### A co-estimated evolutionary timeline of nodulating *Bradyrhizobium* and legumes based on mitochondrial endosymbiosis

To determine whether nodulation arose contemporaneously with legumes, we estimated divergence time using molecular dating. Because bacterial fossils are scarce, we applied a mitochondrial endosymbiosis-based strategy (Shih and Matzke 2013; Wang and Luo 2021) that jointly infers the evolutionary timeline of alphaproteobacterial lineages, including *Bradyrhizobium*, and eukaryotes (including legumes), by leveraging 11 calibrations within the mitochondrial clade (Data S3; Fig. S6) under the scenario that mitochondria originated from a lineage closely related to Alphaproteobacteria (Andersson et al. 1998; Wang and Wu 2015; Martijn et al. 2018). Relaxed molecular clock dating was performed based on 41 conserved Alphaproteobacteria mitochondria-related genes, comprising 24 mitochondria-encoded genes and 17 nuclear-encoded ones of mitochondrial origin (Table S1; see also Methods).

To enable molecular clock analysis in MCMCtree under the site-heterogeneous substitution model LG+C60+G, which is not natively implemented in the software, we incorporated two recently developed methods: phyloHessian and an enhanced version of bs_inBV (see Methods and Note S1). Complex mixture substitution models like LG+C60+G or CAT-GTR are particularly suitable for deep-time evolutionary inference because of its improved accommodation of site-heterogeneous substitution patterns (Schrempf et al. 2020; Pisani et al. 2026). Previous studies have demonstrated the importance of using mixture models in molecular dating, but have focused primarily on deep-branching or uniformly sampled phylogenies (Wang and Meade 2026). We further demonstrated, by simulations matching the diversification pattern of our own dataset, that mixture models consistently outperform their non-mixture counterparts (LG+G in our test) in phylogenies resembling the case of *Bradyrhizobium* analyzed in the present study, where the lineages of interest (*Bradyrhizobium*) are substantially younger than the root and most branching events occur recently (Figs. S7-S9). Furthermore, phyloHessian and bs_inBV performed comparably in terms of divergence time estimates compared to the simulated values under the mixture model, despite using different approaches to approximate the Hessian matrix used in approximate-likelihood dating (dos Reis and Yang 2011): phyloHessian estimates the Hessian by finite differences, whereas bs_inBV approximates it as the inverse of the bootstrap covariance matrix.

We selected 65 representative *Bradyrhizobium* genomes together with 32 alphaproteobacterial genomes and 36 (eukaryotic) mitochondrial lineages for dating analysis under both a single and two partitions of the alignment (Figs. 2, S10). Under the focal calibration strategy, the last common ancestor (LCA) of *Bradyrhizobium* was dated to 212 million years ago (Ma) (95% HPD [Highest Posterior Density] interval, 255-171 Ma), followed by rapid diversification into five supergroups with nodulating members (*B. japonicum*, *B. elkanii*, *B. jicamae*, Kakadu, and PB [Photosynthetic *Bradyrhizobium*]) around 186 Ma (95% HPD interval 224-153 Ma) (the two-partition strategy showed a better model fit as shown in Table S2, therefore we report its time estimates unless otherwise mentioned; both strategies gave similar ages). The crown group of nodulating lineages within the *B. japonicum* supergroup emerged around 88 Ma (95% HPD, 106-72 Ma), and its total group emerged around 97 Ma (95% HPD, 116-80 Ma). This age overlapped with the estimated origin of the *nod*-free but nodulating supergroup (100 Ma; 95% HPD interval 124-76 Ma) and the origin of nodulating legumes (91 Ma; 95% HPD interval 113-77 Ma). Other Nod factor*-*dependent nodulating lineages arose later: *B. jicamae* at 62 Ma (95% HPD interval: 81-45 Ma), *B. elkanii*-1 at 45 Ma (95% HPD interval: 62-30 Ma), and *B. elkanii*-2 at 42 Ma (95% HPD interval: 56-29 Ma) (Fig. 2). This suggests that nodulation first originated in the *B. japonicum* supergroup and subsequently spread across the genus through HGT, although the detailed sequence of HGT events is not clearly solved (Fig. 1b).

**Figure 2.**
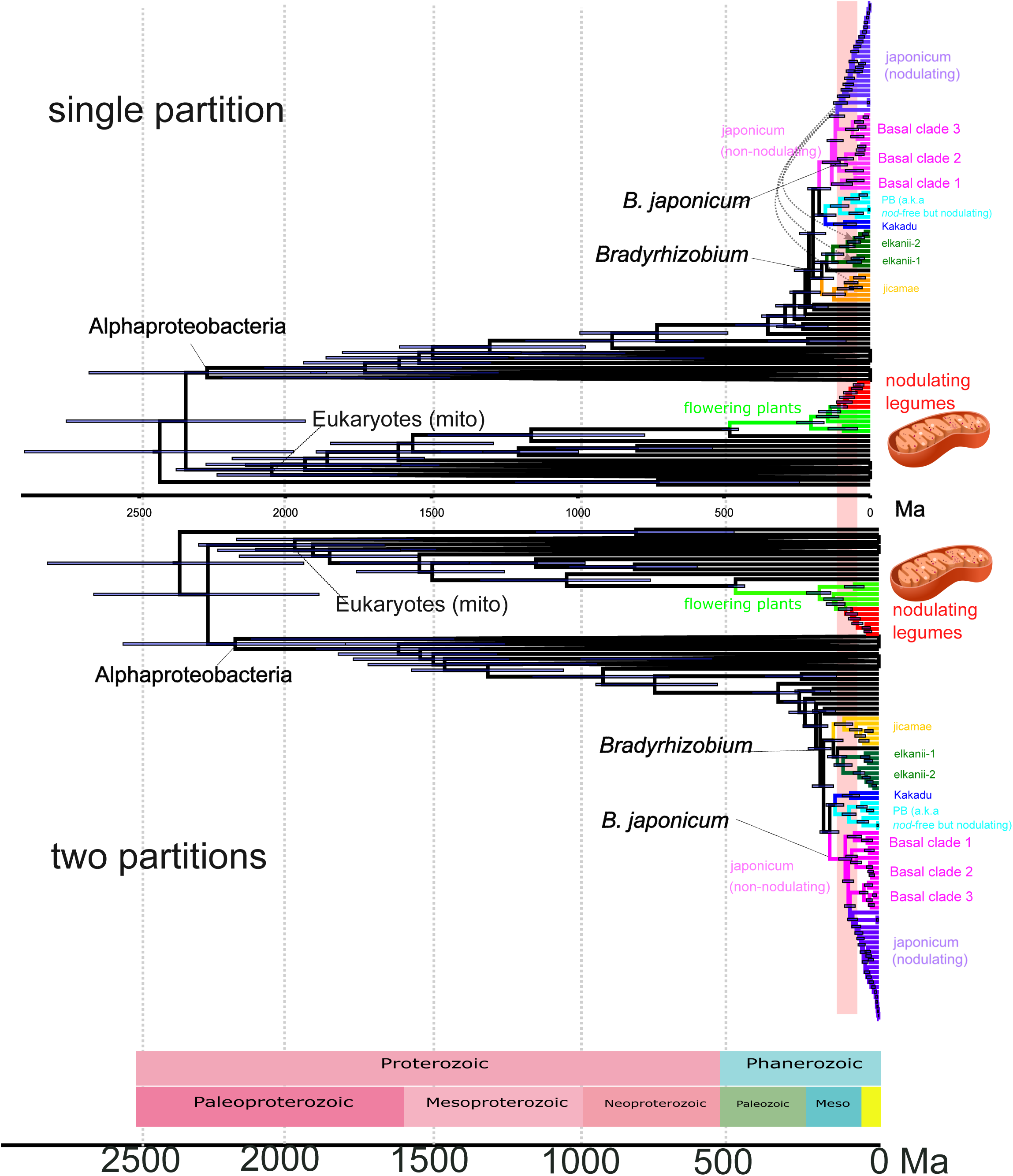
Co-estimating the divergence time of *Bradyrhizobium* and legume host using the mitochondrial endosymbiosis-based strategy under the substitution model LG+C60+G. The timeline is estimated with the *focal* calibration scheme using a single partition (top) or two partitions (bottom). Calibrations are imposed on eukaryotic lineages placed as a sister to Alphaproteobacteria (see Fig. S6 for fossil calibrations). Bars on the node denote the 95% HPD interval of estimated dates. The potential divergence time of nodulating legumes is indicated by a red vertical bar. The image of mitochondrion is credited to Kelvin Song and distributed under Creative Commons Attribution-Share Alike 3.0 (https://creativecommons.org/licenses/by-sa/3.0/).

### Including basal non-nodulating lineages greatly improves time estimates and supports coevolution with legumes

When we removed the three basal non-nodulating clades from the analysis, the estimated age of the LCA of nodulating *B. japonicum* became older than the origin of nodulating legumes by more than 20% with the *focal* calibration strategy under the substitution model LG+C60+G (Fig. 3a). To assess the impact of fossil calibrations on time estimates, we repeated the analyses one by one under a broad range of alternative time-constraint schemes (see Data S3 and Note S2 for a full description). Under the calibration strategies *Cauchy, legume100, legume125 and root4500*, the posterior age estimates of lineages of *Bradyrhizobium* and legumes did not change much (Fig. 3a). In other instances, however, the application of a more disputable and less conservative time calibration for animals and/or red algae (as illustrated by the calibration schemes *Bengtson2017*, *Turner2019* and their combination *Bengtson_Turner* in Fig. 3a) substantially shifted absolute posterior age estimates toward the past (Fig. 3). Nonetheless, in our mitochondria-based divergence co-estimation framework, the absolute divergence times of different lineages also shifted concertedly with each calibration scheme, meaning that the relative order of divergence between lineages did not change: the LCA of nodulating *B. japonicum* remained concurrent with that of legumes when the basal non-nodulating clades were included, but older when they were excluded (Fig. 3a). For example, including basal non-nodulating *B. japonicum* lineages under *Bengtson_Turner* increased the posterior mean divergence time between nodulating *B. japonicum* and legumes to 107 and 109 Ma, respectively, an increase of approximately 20% relative to the focal scheme (the leftmost element in the box), while excluding them changed estimates to 130 and 111 Ma (Fig. 3a, two partitions). In other words, although alternative calibrations altered the absolute divergence-time estimates, it did not change the conclusion that nodulating *B. japonicum* and nodulating legumes diverged contemporarily only when the basal non-symbiotic *B. japonicum* strains were included.

**Figure 3.**
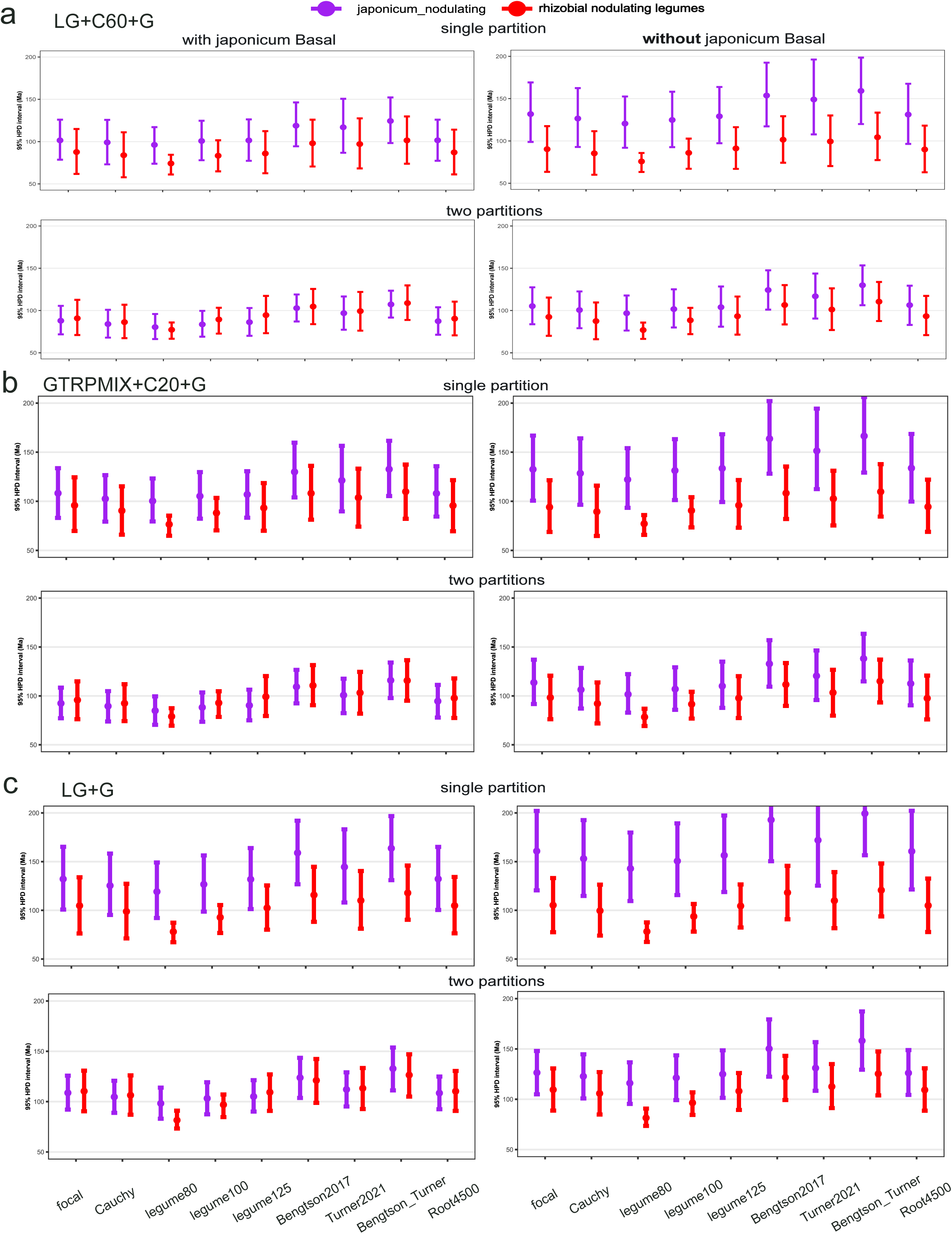
Comparison across calibration schemes of the 95% HPD interval for the estimated divergence time of the crown group of nodulating *B. japonicum*, with basal non-nodulating lineages included (left) or excluded (right), alongside the corresponding estimates for the crown group of nodulating legumes. Three substitution models are tested: compositionally site-heterogeneous model LG+C60+G (a), *Bradyrhizobium*-specific compositionally site-heterogeneous model GTRpmix+C20+G (b), and a traditional site-homogeneous model LG+G (c). Divergence times are analyzed using MCMCtree’s approximate likelihood method with the Hessian matrix estimated under different substitution models using phyloHessian (LG+C60+G and LG+G) and bs_inBV (GTRmpix+C20+G). The centre corresponds to the posterior mean of the estimated divergence time. Both a single partition and two partitions are tested. Nine calibrating schemes are used. *focal*: the focal scheme (see Note S2 and Data S3 for details of each calibration scheme); *Cauchy*, those internal nodes without a proper maximum time bound calibrated as a Cauchy distribution; *legume80, legume100, legume125*: the maximum time bound set to 80, 100, and 125 Ma for nodulating legumes respectively; *Bengtson2017*: the minimum bound of crown-group red algae set as 1600 Ma; *Turner2019*: the minimum time bound of the crown-group Metazoa (total group of sponges) set as 890 Ma; *Bengtson_Turner*: a combination of *Bengtson2017* and *Turner2019*; *root45*: root age set as 4500 Ma instead of 3000 Ma.

In addition, molecular clock dating analyses were performed under alternative mixture models to check the robustness of time estimates to substitution models. These include LG+C20+G (Fig. S11a), which uses 20 rather than 60 empirical equilibrium-frequency vectors; LG+C60+R+I (Fig. S11a), which incorporates a FreeRate model (+R) of among-site rate variation and an invariant-site component (+I); and GTRpmix+C20+G (Fig. 3b), a data-specific mixture substitution model. All of them recovered the same divergence order as the focal LG+C60+G model. The poorly fitting site-homogeneous LG+G model (Table S2) under a single-partition analysis, however, inferred that nodulating *B. japonicum* originated earlier than legumes even if those basal non-symbiotic lineages were included with some calibration strategies, producing an evolutionarily less plausible event (Fig. 3c). The above pattern was further supported by analysis with an alternative set of genes (*gene53* in Fig. S12). We further used a more conservative calibration strategy by moving the calibration on the crown-group red algae in each calibration strategy to the total-group red algae, and the patterns remained the same (Fig. S13; see Note S2.2.7 for a detailed discussion).

## Discussion

Symbiosis has transformed the evolution of life by enabling organisms to acquire new functions and exploit novel ecological niches. Despite its broad evolutionary importance, how and when free-living bacteria transition into mutualistic symbionts remains poorly understood. The *Bradyrhizobium*-legume symbiosis offers a powerful system to address this question because the genus encompasses both free-living nitrogen-fixing lineages and legume-nodulating symbionts (Kulkarni et al. 2015; Jones et al. 2016; Tao et al. 2021; Ling et al. 2026). By combining broad sampling of non-symbiotic lineages with improved molecular dating under a mitochondrial endosymbiosis-based framework, the present study provides a temporal resolution of how rhizobial symbiosis evolved. The mitochondrial endosymbiosis-based framework jointly dates host and symbiont lineages within a single phylogenetic analysis (Fig. 2). A desirable property of this approach is that alternative fossil calibrations shift absolute ages in a concerted manner while preserving the relative divergence order between lineages (Wang and Luo 2021). Clearly, more recent calibrations change divergence time estimates toward the present, whereas older calibrations push them deeper into the past. Hence, when hosts and symbionts are analyzed together in a single analysis, changes in the calibrations are expected to affect only their absolute divergence times, but not the relative ordering. This matters since comparisons of divergence time estimates across studies may suggest conflicting evolutionary timelines which, however, may simply reflect differences in the calibrations applied to legumes and rhizobia, thereby leading to a misleading comparison.

The identification of three basal non-nodulating clades within the *B. japonicum* supergroup provides important evolutionary anchors for this framework. This substantially reduces the possibility that there are more basal unsampled symbiotic lineages. Using these anchors with the best-fitting mixture substitution models (Table S2), the analyses show that nodulating *Bradyrhizobium* originated contemporaneously with nodulating legumes around 90 Ma, rather than well before or after them. Excluding these basal non-symbiotic lineages distorts the temporal signal and falsely implies that rhizobia predated legumes (Figs. 3, S11-S13). These patterns were robust across alternative mixture substitution models. However, when the homogeneous LG+G model was applied with a single partition, the inferred origin of nodulating *Bradyrhizobium* still predated that of nodulating legumes, even with the basal non-symbiotic lineages. Moreover, under the focal calibration schemes, mixture models produced a time estimate of legume divergence closer to previous estimates, which is around 60 to 110 Ma (Hohmann et al. 2015; Foster et al. 2017; Nieto-Blázquez et al. 2017; Zhao et al. 2021), than homogeneous models (Fig. 3). This indicates that a broader taxon sampling alone may be insufficient to recover the biologically plausible evolutionary scenario, and should therefore be combined with mixture models as a best practice. We applied two approaches for applying complex mixture substitution models in dating under the approximate likelihood method via Hessian estimation, extending its calculation previously designed under only homogeneous models (Thorne et al. 1998; dos Reis and Yang 2011): (1) a direct calculation with phyloHessian (Wang and Meade 2026), and (2) a bootstrap-based approximation via bs_inBV (Wang and Luo 2025). When the mixture substitution model is available in phyloHessian or IQ2MC (Demotte et al. 2025), the first approach is preferred for its higher accuracy and speed; otherwise, bs_inBV provides a good alternative for data-specific models such as GTRpmix, as long as the phylogenetic likelihood can be correctly calculated.

Now, three temporal scenarios for the origin of rhizobial nodulation can be distinguished: nodulation capacity may have originated before legumes; nodulating *Bradyrhizobium* and nodulating legumes may have arisen contemporaneously, consistent with reciprocal co-evolution; or rhizobia may have acquired nodulation only after legumes had diversified, implying one-sided bacterial adaptation. The results favor contemporaneous emergence as the most parsimonious interpretation. The first scenario receives the least support, not only because it is inconsistent with the molecular dating results, but also because an origin of nodulating *Bradyrhizobium* well before legumes would predict a broader non-legume host range. Rhizobial nodulation is known only from nodulating legumes (Caesalpinioideae and Papilionoideae) and the small rosid genus *Parasponia* (Doyle 2016; Remigi et al. 2016). Nodulating plants comprise two major symbiotic types: actinorhizal nodulation with *Frankia*, found across multiple lineages of the nitrogen-fixing clade consisting of Fabales (including legumes), Rosales (including *Parasponia*), Fagales, and Cucurbitales; and rhizobial nodulation with rhizobia, found mainly in legumes and independently in *Parasponia* (Doyle et al. 2025). Several recent studies suggest that plant nodulation originated only once, possibly as actinorhizal nodulation, in the LCA of Fabales, Rosales, Fagales, and Cucurbitales (Griesmann et al. 2018; van Velzen et al. 2018; Bu et al. 2020). The results presented here imply that rhizobial nodulation was derived from this ancestral actinorhizal system in legumes most likely around 90-80 Ma.

Several caveats should be considered when interpreting these findings. Absolute ages are sensitive to calibration choices and clock model assumptions, even when relative node order is stable. Taxon sampling is substantially improved but still incomplete, with sparse Kakadu representation and uneven geographic coverage, which may affect estimated ages and inferred transition patterns. Future genomic analyses of more basal lineages could reveal the transitions from a free-living ancestor to nodulating species.

Despite these caveats, the ancient origin of *Bradyrhizobium* indicates that it was predominantly free-living during the first half of its evolutionary history before establishing symbiosis with legumes approximately 90 Ma, coincident with the emergence of nodulating legumes. These findings support a reciprocal model in which legume diversification created ecological opportunities that likely favored the evolution of symbiotic traits in *Bradyrhizobium*, while the resulting associations may have in turn influenced host diversification and ecological expansion. This framework might also apply to other host-microbe associations involving transitions from free-living to symbiotic lifestyles although future work is needed to explore this possibility.

## Methods

### Bacterial sampling and isolation

To increase the diversity of non-leguminous plant-associated *Bradyrhizobium*, we collected non-legume plants, including *Oryza sativa* subsp. *indica* and *japonica*, *Camphora officinarum* and *Houttuynia cordata*, as well as nearby soils (0-5 cm depth) from Hunan (27.948 °N, 113.221 °E), China, in 2021. Samples were individually sealed in sterile bags, kept on ice during transport, and processed immediately upon arrival in the lab. Each sample was separated into root, rhizosphere, and soil compartments for bacterial isolation (detailed in the following paragraph). Basic soil properties were also measured to support subsequent biological interpretations; these data are provided in Data S1.

A series of tenfold dilutions of rhizosphere and soil samples were prepared and then inoculated (100 µL) into the modified N-free arabinose-gluconate media (1.0 g DL-arabinose, 1.0 g sodium gluconate, 1.0 g yeast extract, 2 mL KH_2_PO_4_ solution (110 g/L), 4 mL Na_2_SO_4_ solution (62.5 g/L), 1 mL MgSO_4_•7H_2_O solution (180 g/L), 1 mL CaCl_2_ solution (13 g/L), 1 mL FeCl_3_•6H_2_O solution (6.7 g/L) and 15 g agar were added to 1 L Milli-Q water. The pH was adjusted to 6.6 with KOH, after which the medium was autoclaved at 121 ℃ for 15 to 30 minutes (Tao et al. 2021) and incubated at 28 ℃ for at least 7 days to allow *Bradyrhizobium* to grow. Root sterilization was performed with 75% ethanol and 5% sodium hypochlorite according to the study (Mbai et al. 2013). The roots were washed and ground in a sterilized mortar after sterilization. Fivefold and tenfold dilutions of each grounded root sample were made with PBS buffer (pH 6.8) and then followed the same procedure as soil samples. The identification procedure of each isolate was performed according to (Tao et al. 2021).

### Reconstruction of phylogenomic and *nod*-gene tree

*Bradyrhizobium* genomes were downloaded from NCBI GenBank (last accessed in June 2022) with additional genomes sequenced in two recent studies (Ling et al. 2025; Ling et al. 2026). Phylogenomic analysis was based on 123 single-copy genes across *Bradyrhizobium* identified in a previous study, or 891 single-copy genes within the *B. japonicum* supergroup identified by OrthoFinder v3.0.1 (Emms and Kelly 2019). Gene trees of *nod* genes were built with *nodABCIJ* or *nodABC*. Protein sequences were aligned using MAFFT v7.222 (Katoh and Standley 2013), and was further trimmed using TrimAl v1.4 (Capella-Gutiérrez et al. 2009) with the settings “-automated1 -resoverlap 0.55 -seqoverlap 60” for phylogenomic datasets. For building *nod*-gene trees, which contained only a few genes, alignments were not trimmed. Phylogenies were built using IQ-Tree v3.0.1 (Wong et al. 2026) preferably with mixture substation models Cxx (where xx represents 10-60 pre-defined mixture components) (Le et al. 2008) under a PMSF approximation (Wang et al. 2018), specified by the parameter “-m LG+Cxx+G -f guide_tree -mwopt” where the guide tree to calculate the PMSF was constructed by LG4M (Le et al. 2012). Alternatively, trees were constructed using the site-homogeneous model LG+G (*Bradyrhizobium* phylogenomic tree) or the best-fitting substitution models (*nod*-gene tree) for each partition (“-s alignment -spp partition -m MFP - mset LG -mrate E,G,I,G+I”). Branch support was assessed by 1000 ultrafast bootstraps implemented in IQ-Tree (Hoang et al. 2018). Visualization and operation of the phylogenies were conducted with iTOL v7 (Letunic and Bork 2024), FigTree v1.4.5 (Rambaut 2010) and Newick Utilities (Junier and Zdobnov 2010).

### Collection of genes for molecular clock dating

The mitochondrial endosymbiosis-based strategy places mitochondria as sister to Alphaproteobacteria, uses eukaryotic fossils as time calibrations to date Alphaproteobacteria evolution (Wang and Luo 2021), and has been applied to more distantly related bacteria (Mahendrarajah et al. 2023; Liao et al. 2024; Davín et al. 2025; Wang and Luo 2025; Kay et al. 2026). Our previous study (Wang and Luo 2021) identified 46 conserved genes shared between Alphaproteobacteria and mitochondria, comprising 24 mitochondrially encoded genes and 22 nuclear-encoded genes of mitochondrial origin. After removing those potentially encoded by both the mitochondrial and nuclear genomes (duplicates), we retrieved 41 genes (24 mito-encoded and 17 nuclear-encoded) genes which are the default set of genes if not otherwise indicated (*gene41*). These genes were originally identified in previous studies (Wang and Wu 2015; Martijn et al. 2018) including 24 genes encoded by the mitochondrial genome and 29 genes encoded by the nuclear genome (the full gene set: *gene53*; see also Table S1).

### Genomes included in molecular dating

Genomes used in the dating analysis consisted of 65 *Bradyrhizobium* selected from the phylogenomic tree using TreeCluster v1.0.4 (Balaban et al. 2019) with “-k 0.4” to group genomes into clusters from which one representative was randomly selected per cluster (we chose those isolated from the present study if more than one genome was included in the cluster), and 32 alphaproteobacterial genomes selected from and 36 eukaryotic lineages used in our previous study (Wang and Luo 2021). We additionally included six nodulating legumes (*Medicago truncatula*, *Trifolium pratense*, *Pisum sativum*, *Lotus japonicus*, *Glycine max*, and *Lupinus albus*), one early-branching non-nodulating legume (*Cercis canadensis*), and two non-legume flowering plants (see Table S3 for details). We considered the LCA of *Clycine* (Caesalpinioideae) and *Lupinus* (Papilionoideae) as the origin of nodulating legumes (Fig. S6, Node 13). The combined clade of Caesalpinioideae and Papilionoideae includes all legumes that can form root nodules with rhizobia, although a small proportion (∼30%) have lost this ability, consistent with the model of a single origin followed by multiple independent losses (Zhao et al. 2021).

### Molecular clock dating analysis under mixture substitution model

We used MCMCtree implemented in PAML v4.10.0 (Yang 2007) for molecular dating. The time unit was set to 100 Ma. The prior on divergence times was constructed with fossil calibrations and a birth-death process (Yang and Rannala 1997). The parameters of the birth-death process were set as: birth rate = 1, death rate = 1, and the taxon sampling proportion = 0, which specifies a flat time prior. The independent rate (IR) model, which showed significantly much better fit compared to the autocorrelated-rates (AR) model based on Bayesian model selection with the approximate likelihood method by mcmc3r (dos Reis et al. 2018; Panchaksaram et al. 2025), was used (Table S4). The substitution rate for each gene partition was assigned a Dirichlet-gamma prior (“rgene_gamma = 1 50 1”), corresponding to a mean rate of 0.02 substitutions/100 Myr. We set the burn-in, sampling frequency, and iteration number as *2* × *10^5^*, 100, and *3* × *10^4^*, respectively to ensure that the effective sample size (ESS) calculated by R package coda (Plummer et al. 2006) was larger than 200. To make sure that the sequence data was informative, we compared the posterior ages and effective priors by setting *usedata* = 0 in the control file *mcmctree.ctl*. Following (Wang and Meade 2026), the sequence alignment was analyzed either as a single partition or as two partitions defined by a two-component Gaussian mixture model fitted to the logarithm of the estimated evolutionary rate of each gene.

MCMCtree implements only profile homogeneous models like LG and WAG combined with Gamma across-site relative rate heterogeneity. To enable the use of the mixture substitution model in dating with MCMCtree, we applied our recently developed software phyloHessian v0.4.0 (Wang and Meade 2026). The software uses a finite-difference method (Seo et al. 2004; dos Reis and Yang 2011) to compute the gradient and Hessian, i.e., the first- and second-order partial derivatives of the log-likelihood function with respect to branch lengths, evaluated at the maximum-likelihood branch length estimates, under not only site-homogeneous model available in MCMCtree but a wide range of substitution models (Demotte et al. 2025), particularly mixture substitutions such as Cxx (Le et al. 2008) and the PMSF approximation (Wang et al. 2018). These derivatives were then employed in MCMCtree’s approximate likelihood framework to approximate the log-likelihood surface during divergence-time inference (Thorne et al. 1998; dos Reis and Yang 2011). We further improved bs_inBV which was originally developed in our previous study (Wang and Luo 2025) by fixing the nuisance parameters and by better dealing with zero-length branches in Hessian approximation (Note S1). It uses a bootstrap approach to approximating the Hessian. It was applied when the substitution model was not implemented in phyloHessian (see Note S1.1). Specifically, we estimated a new, dataset-specific exchangeability matrix using IQ-Tree (“-m GTR20+C20+G --link-exchange --init-exchange LG -me 0.999”) with the GTRpmix profile-mixture model (Baños et al. 2024). We then used bs_inBV to run MCMCtree with the bootstrap-estimated Hessian derived from the substitution model GTRpmix+C20+G.

## Data and code Availability

Genome sequences of newly isolated strains are deposited at NCBI Genbank. Updated bs_inBV which allows for the bootstrap approach to using MCMCtree’s approximate likelihood molecular dating under diverse substitution models is available at https://github.com/evolbeginner/bs_inBV. Data and code are available at FigShare https://figshare.com/s/fec450d9b9de34cf8998. The genomic sequences and raw reads of the 10 newly sequenced *Bradyrhizobium* isolates are available at NCBI BioProject ID: PRJNA1469367 (https://dataview.ncbi.nlm.nih.gov/object/PRJNA1469367?reviewer=sf3vtkuqjjqotqr1n24cioi4pm). The interactive interface for chronogram visualization and manipulation (chronogram_viewer) is available at https://sishuowang.github.io/chronogram_viewer/.

## Acknowledgments

We thank Luo lab member Jinjin Tao for the discussion, Xinyu Ma, Yan Zuo, and Xing Yang for sample collection in China. We also thank Hector Baños and Piyumal Demotte for advice on phylogenetic analysis, and Jun Huang for statistical analysis. This work is supported by the Hong Kong Research Grants Council General Research Fund (Grant No. 14116922 [sequencing, phylogenomics, molecular dating] and 14118725 [molecular clock model development]), and the Natural Science Foundation of China (Grant No. 32400493).

