## Supplementary material for "Molecular clock dating under mixture substitution models indicates contemporaneous emergence of *Bradyrhizobium* nodulation and legumes anchored by non-symbiotic relatives": Tables S1-S4, Figs. S1-S13, Notes S1-S2

September 5, 2026

**Supplementary Figures** **8**

**Supplementary Notes** **22**

**S1 Additional notes on molecular clock dating** **22**

S1.1 Improved bootstrap-based approach for using mixture substitution models in MCMCtree 22

#### Supplementary Table

Table S1: Gene sets used in molecular dating. The full set comprises 53 genes (*gene53*), of which 41 (*gene41*) carefully selected ones were used in the main analyses. Gene origin is indicated as "Mito" (mitochondrially encoded) or "Nuclear" (nuclear-encoded genes of a mitochondrial evolutionary origin). Duplicate gene pairs are indicated in the "Note" column. The gene set *gene53* is used only in the analysis displayed in fig. S12.

| **No.** | **Gene** | **Encoded by** | ***gene41*** | **Note** |
|---|---|---|---|---|
| 1 | AFG1 | Nuclear | ✓ | |
| 2 | apaG | Nuclear | | |
| 3 | bioC | Nuclear | ✓ | |
| 4 | clpB | Nuclear | | |
| 5 | clpP | Nuclear | | |
| 6 | cox11 | Nuclear | ✓ | |
| 7 | dnaK | Nuclear | ✓ | |
| 8 | duf185 | Nuclear | ✓ | |
| 9 | engA | Nuclear | ✓ | |
| 10 | erpA | Nuclear | ✓ | |
| 11 | gidA | Nuclear | ✓ | |
| 12 | groEL | Nuclear | ✓ | |
| 13 | grpE | Nuclear | ✓ | |
| 14 | hemN | Nuclear | | |
| 15 | hesB | Nuclear | ✓ | |
| 16 | hslV | Nuclear | | |
| 17 | ksgA | Nuclear | | |
| 18 | mito-MitoCOG0001 | Mito | ✓ | |
| 19 | mito-MitoCOG0003 | Mito | ✓ | |
| 20 | mito-MitoCOG0004 | Mito | ✓ | |
| 21 | mito-MitoCOG0005 | Mito | ✓ | |
| 22 | mito-MitoCOG0008 | Mito | ✓ | |
| 23 | mito-MitoCOG0009 | Mito | ✓ | |
| 24 | mito-MitoCOG0010 | Mito | ✓ | |
| 25 | mito-MitoCOG0011 | Mito | ✓ | |
| 26 | mito-MitoCOG0012 | Mito | ✓ | |
| 27 | mito-MitoCOG0027 | Mito | ✓ | |
| 28 | mito-MitoCOG0030 | Mito | ✓ | |
| 29 | mito-MitoCOG0031 | Mito | ✓ | Duplicate of nuoD |
| 30 | mito-MitoCOG0039 | Mito | ✓ | |
| 31 | mito-MitoCOG0040 | Mito | ✓ | |
| 32 | mito-MitoCOG0043 | Mito | ✓ | Duplicate of nuoC |
| 33 | mito-MitoCOG0052 | Mito | ✓ | Duplicate of sdhB |
| 34 | mito-MitoCOG0053 | Mito | ✓ | |
| 35 | mito-MitoCOG0055 | Mito | ✓ | |
| 36 | mito-MitoCOG0059 | Mito | ✓ | |
| 37 | mito-MitoCOG0060 | Mito | ✓ | |
| 38 | mito-MitoCOG0066 | Mito | ✓ | Duplicate of nuoG |
| 39 | mito-MitoCOG0067 | Mito | ✓ | |

| No. | Gene | Encoded by | *gene41* | Note |
|---|---|---|---|---|
| 40 | mito-MitoCOG0071 | Mito | ✓ | Duplicate of nuoI |
| 41 | mito-MitoCOG0133 | Mito | ✓ | |
| 42 | mraW | Nuclear | | |
| 43 | nuoC | Nuclear | | Duplicate of mito-MitoCOG0043 |
| 44 | nuoD | Nuclear | | Duplicate of mito-MitoCOG0031 |
| 45 | nuoF | Nuclear | ✓ | |
| 46 | nuoG | Nuclear | | Duplicate of mito-MitoCOG0066 |
| 47 | nuoI | Nuclear | | Duplicate of mito-MitoCOG0071 |
| 48 | petA | Nuclear | ✓ | |
| 49 | rpl3 | Nuclear | ✓ | |
| 50 | sdhB | Nuclear | | Duplicate of mito-MitoCOG0052 |
| 51 | sucD | Nuclear | ✓ | |
| 52 | trmE | Nuclear | ✓ | |
| 53 | ybjS | Nuclear | ✓ | |

Table S2: AIC (lower value indicates a better fit) of different substitution models under single- and two-partition schemes in phylogenomic reconstruction based on the focal *gene41* gene set for molecular dating. The number of parameters is calculated as $k = B + G + (M - 1)$, where $B$ is the number of branches, $G = 1$ for +G's $\alpha$, and $M$ is the number of equilibrium frequency vectors ($M = 60$ for C60; otherwise 0). In our case, the branch lengths are estimated for an unrooted tree, thus $B = 135 \times 2 - 3 = 267$.

| Scheme | Model | lnL | Parameter number | AIC |
|---|---|---|---|---|
| **Single partition** | | | | |
| Single partition | LG+G | -839657 | 268 | 1679850 |
| Single partition | LG+C60+G | -817866 | 327 | 1636386 |
| **Two partitions: Partition 1** | | | | |
| Partition 1 | LG+G | -250510 | 268 | 501556 |
| Partition 1 | LG+C60+G | -243625 | 327 | 487904 |
| **Two partitions: Partition 2** | | | | |
| Partition 2 | LG+G | -586900 | 268 | 1174336 |
| Partition 2 | LG+C60+G | -567724 | 327 | 1136102 |

Table S3: Genome sources for the eukaryotes used in molecular dating.

| **Taxonomy** | **Species** | **Nuclear genome** | **Mitogenome** |
|---|---|---|---|
| Metazoa | *Homo sapiens* | see Wang and Luo (2021) | |
| Metazoa | *Gallus gallus* | see Wang and Luo (2021) | |
| Metazoa | *Branchiostoma floridae* | see Wang and Luo (2021) | |
| Metazoa | *Amphimedon queenslandica* | see Wang and Luo (2021) | |
| Fungi | *Candida albicans* | see Wang and Luo (2021) | |
| Fungi | *Ustilago maydis* | see Wang and Luo (2021) | |
| Fungi | *Pleurotus ostreatus* | see Wang and Luo (2021) | |
| Fungi | *Spizellomyces punctatus* | see Wang and Luo (2021) | |
| Amoebozoa | *Acanthamoeba castellanii* | see Wang and Luo (2021) | |
| Amoebozoa | *Dictyostelium discoideum* | see Wang and Luo (2021) | |
| Amoebozoa | *Polysphondylium pallidum* | see Wang and Luo (2021) | |
| Archaeplastida | *Arabidopsis thaliana* | see Wang and Luo (2021) | |
| Archaeplastida | *Oryza sativa* | see Wang and Luo (2021) | |
| Archaeplastida | *Physcomitrella patens* | see Wang and Luo (2021) | |
| Archaeplastida | *Ostreococcus tauri* | see Wang and Luo (2021) | |
| Archaeplastida | *Chondrus crispus* | see Wang and Luo (2021) | |
| Archaeplastida | *Porphyra umbilicalis* | see Wang and Luo (2021) | |
| Archaeplastida | *Cyanidioschyzon merolae* | see Wang and Luo (2021) | |
| Archaeplastida | *Cyanophora paradoxa* | see Wang and Luo (2021) | |
| Heterokontophyta | *Phytophthora infestans* | see Wang and Luo (2021) | |
| Heterokontophyta | *Thalassiosira pseudonana* | see Wang and Luo (2021) | |
| Discoba | *Andalucia godoyi* | see Wang and Luo (2021) | |
| SAR | *Symbiodinium minutum* | see Wang and Luo (2021) | |
| SAR | *Paramecium tetraurelia* | see Wang and Luo (2021) | |
| SAR | *Oxytricha trifallax* | see Wang and Luo (2021) | |
| SAR | *Reticulomyxa filosa* | see Wang and Luo (2021) | |
| SAR | *Elphidium margaritaceum* | see Wang and Luo (2021) | |
| Archaeplastida (legumes) | *Lotus japonicus* | PLAZA (Van Bel et al., 2022) | NC_029641.1 |
| Archaeplastida (legumes) | *Glycine max* | PLAZA | JX463295 |
| Archaeplastida (legumes) | *Medicago truncatula* | PLAZA | NC_016743.2 |
| Archaeplastida (legumes) | *Trifolium pratense* | PLAZA | MW448461.1 |
| Archaeplastida (legumes) | *Lupinus albus* | PLAZA | NC_048499.1 |
| Archaeplastida (legumes) | *Pisum sativum* | PLAZA | MN017226 |
| Archaeplastida (basal non-nodulating legumes) | *Cercis canadensis* | PLAZA | CM044352.1 |
| Archaeplastida (monocots) | *Zea mays* | PLAZA | NC_007982 |
| Archaeplastida (eudicots) | *Vitis vinifera* | PLAZA | NC_012119 |

*(Continued from previous page)*

| Taxonomy | Species | Nuclear genome | Mitogenome |
|---|---|---|---|

*Note: Histiona aroides*, *Jakoba bahamiensis*, *Jakoba libera*, *Reclinomonas americana*, *Seculamonas ecuadoriensis*, and *Malawimonas jakobiformis* in (Wang and Luo, 2021) were not used because their nuclear genomes are not yet sequenced.

Table S4: mcmc3r model comparison for gene41 and gene53 datasets. The gene set *gene41* (41 genes) is used in most analyses, while *gene53* is the full gene set (53 genes). Posterior model probabilities are computed by comparing AR vs. IR within each partition scheme.

| Gene dataset | Partition scheme | log ML (AR) | log ML (IR) | $P(\text{AR} \mid D)$ | $P(\text{IR} \mid D)$ |
|---|---|---|---|---|---|
| gene41 | single partition | $-980.2693 \pm 2.430619$ | $-918.6321 \pm 2.274601$ | 0 | 1 |
| gene41 | two partitions | $-1611.444 \pm 2.512731$ | $-1463.971 \pm 1.959463$ | 0 | 1 |
| gene53 | single partition | $-992.8344 \pm 2.393920$ | $-889.4515 \pm 2.400712$ | 0 | 1 |
| gene53 | two partitions | $-1619.846 \pm 2.273871$ | $-1449.006 \pm 2.266895$ | 0 | 1 |

#### Supplementary Data (see the Excel file)

**S1 Information of strains isolated in the present study.**

**S2 *Bradyrhizobium* strains downloaded from GenBank**

**S3 Calibrations in molecular dating**

#### Supplementary Figures

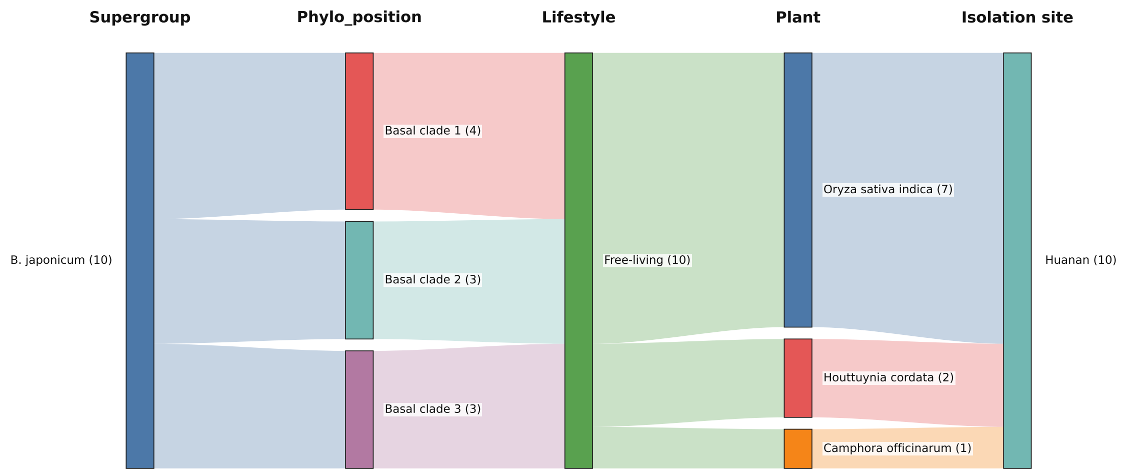

(a)

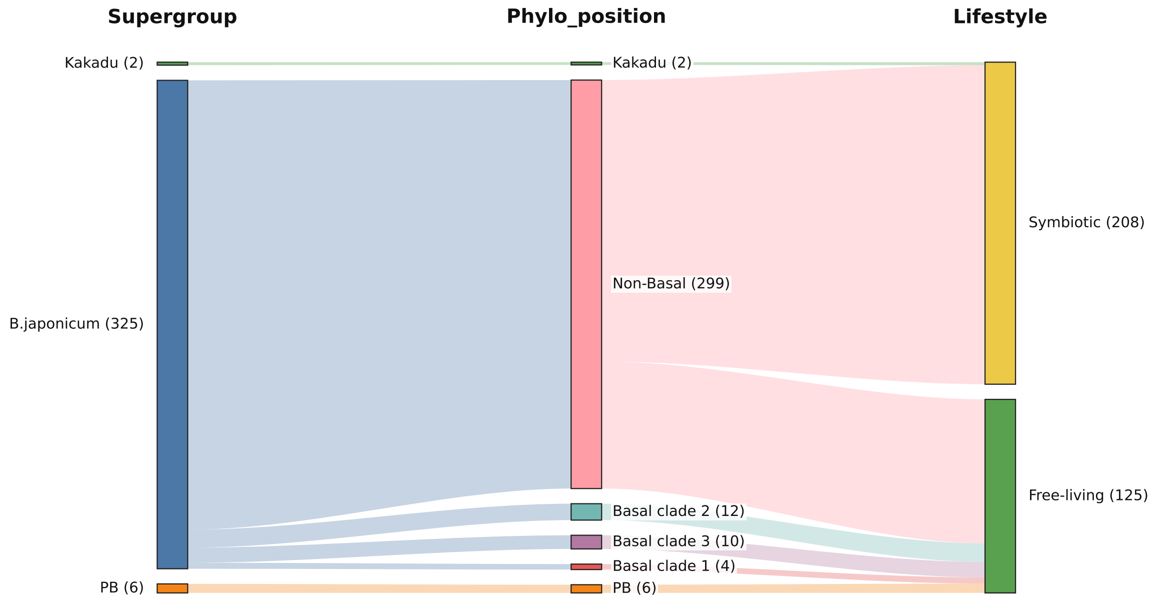

(b)

Figure S1: Sankey diagrams showing the information of *Bradyrhizobium* strains. (a) Strains isolated in the present study. (b) Strains used to reconstruct the phylogenomic tree.

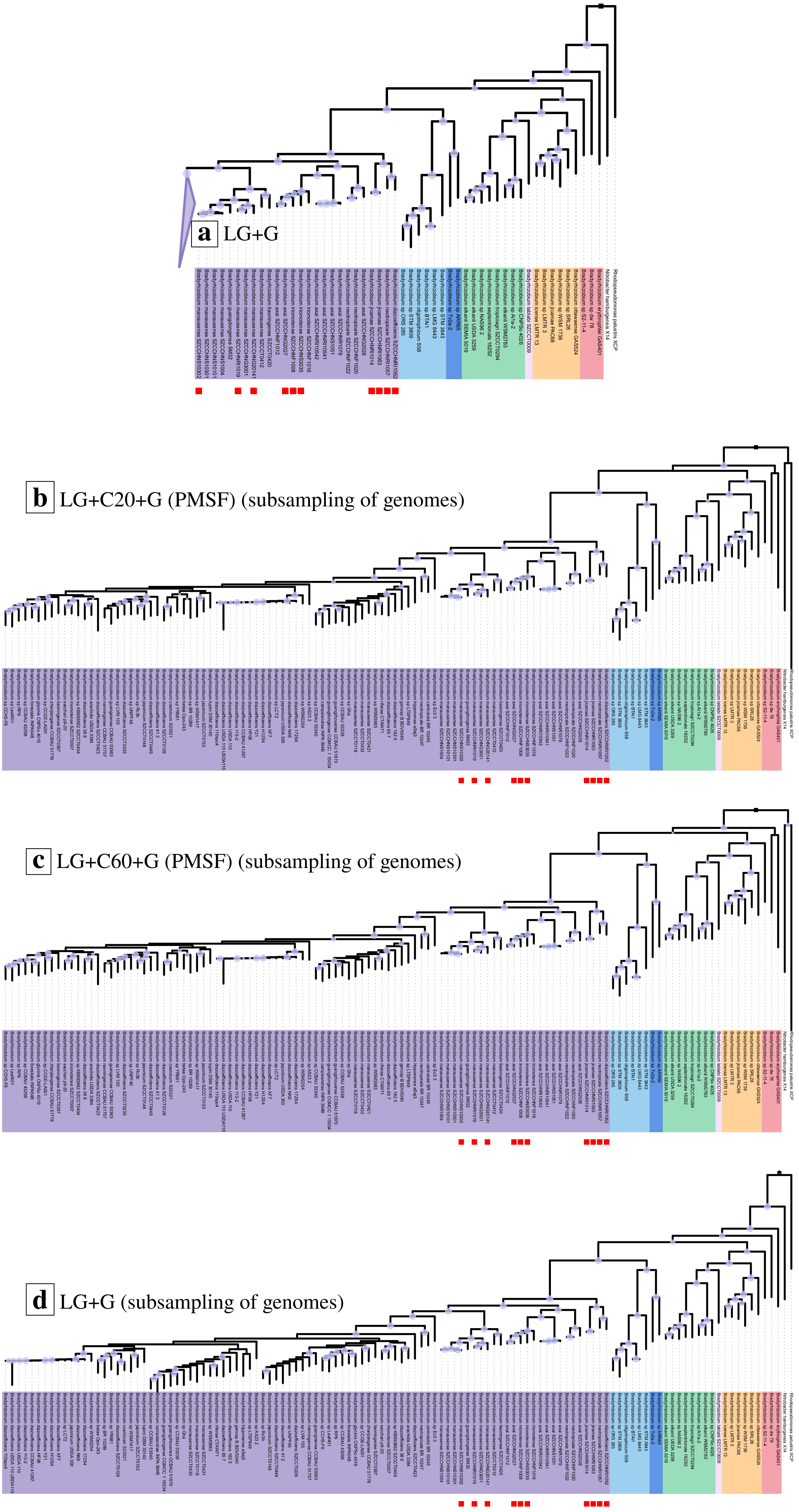

Figure S2: Phylogenomic trees of the *B. japonicum* supergroup using the **123 orthologs (Tao et al., 2021)** conserved across the genus under various substitution models. (a) site-homogeneous model LG+G. (b-d) site-heterogeneous models LG+C20+G (PMSF), LG+C60+G (PMSF), and site-homogeneous model LG+G, under a subsampling of *B. japonicum* genomes. The subsampling of *B. japonicum* genomes (basal non-nodulating lineages excluded) is performed using TreeCluster (Balaban et al., 2019) with a cutoff of phylogenetic depth at 0.04, which groups tips into different clusters and randomly selects one from each, to speed up tree reconstruction under computationally heavy mixture substitution models. Circles on nodes indicate ultrafast bootstrap values of 90 to 100, with circle size increasing from smallest to largest. Red squares at tips indicate strains isolated in the present study.

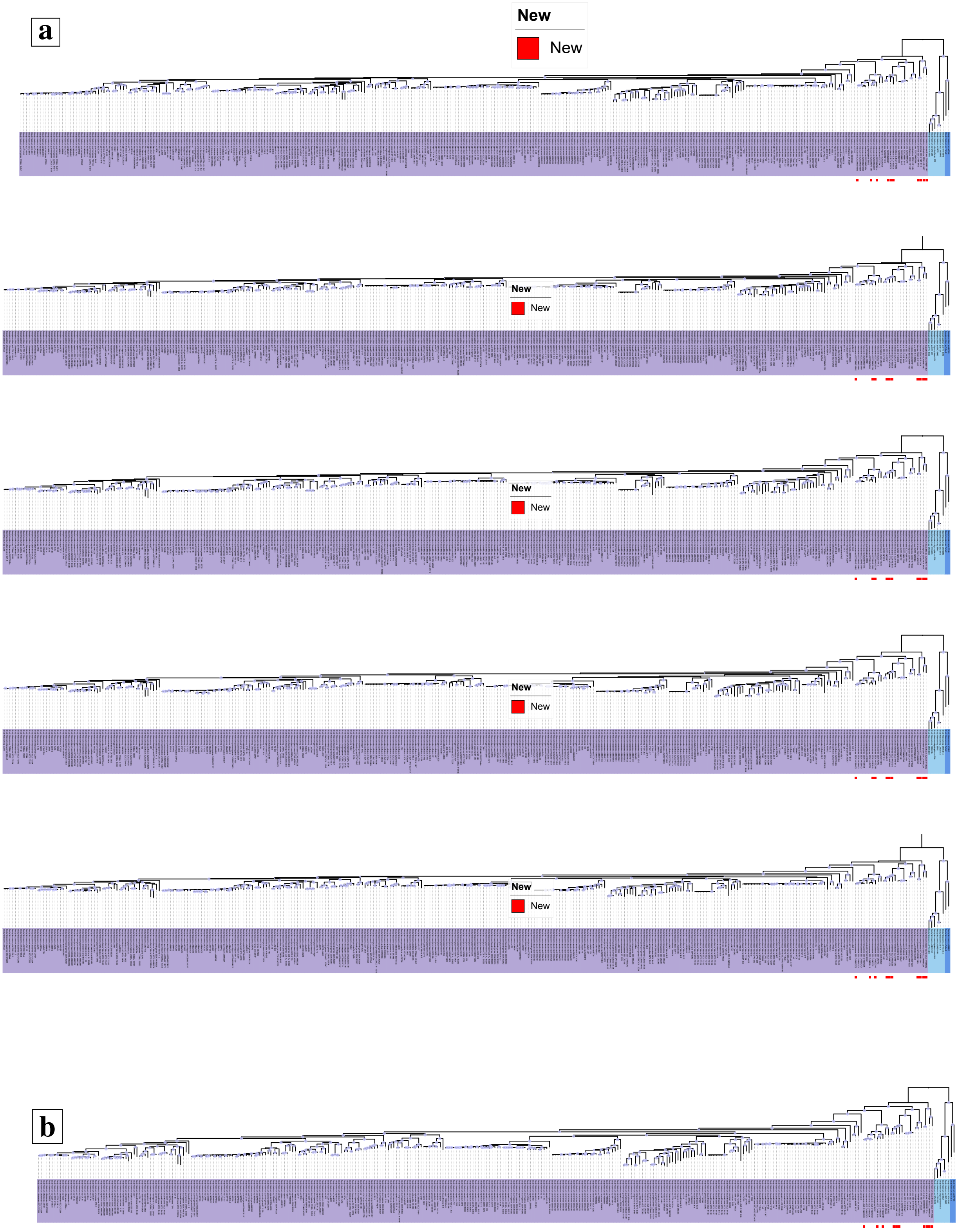

Figure S3: Phylogenomic trees of *B. japonicum* with **891 single-copy genes conserved within the *B. japonicum* supergroup**. (a) site-heterogeneous model LG+C20+G (PMSF) with 100 randomly selected orthologs from the 891 genes, repeated for five times. (b) site-homogeneous model LG+G. A total of 373 (365 *B. japonicum* genomes plus eight from the PB (*nod*-free nodulating) and Kakadu supergroups as the outgroup) genomes are included. Cyan circles on nodes indicate ultrafast bootstrap values of 90-100, with circle size increasing from smallest to largest. Red squares at tips indicate strains isolated in the present study.

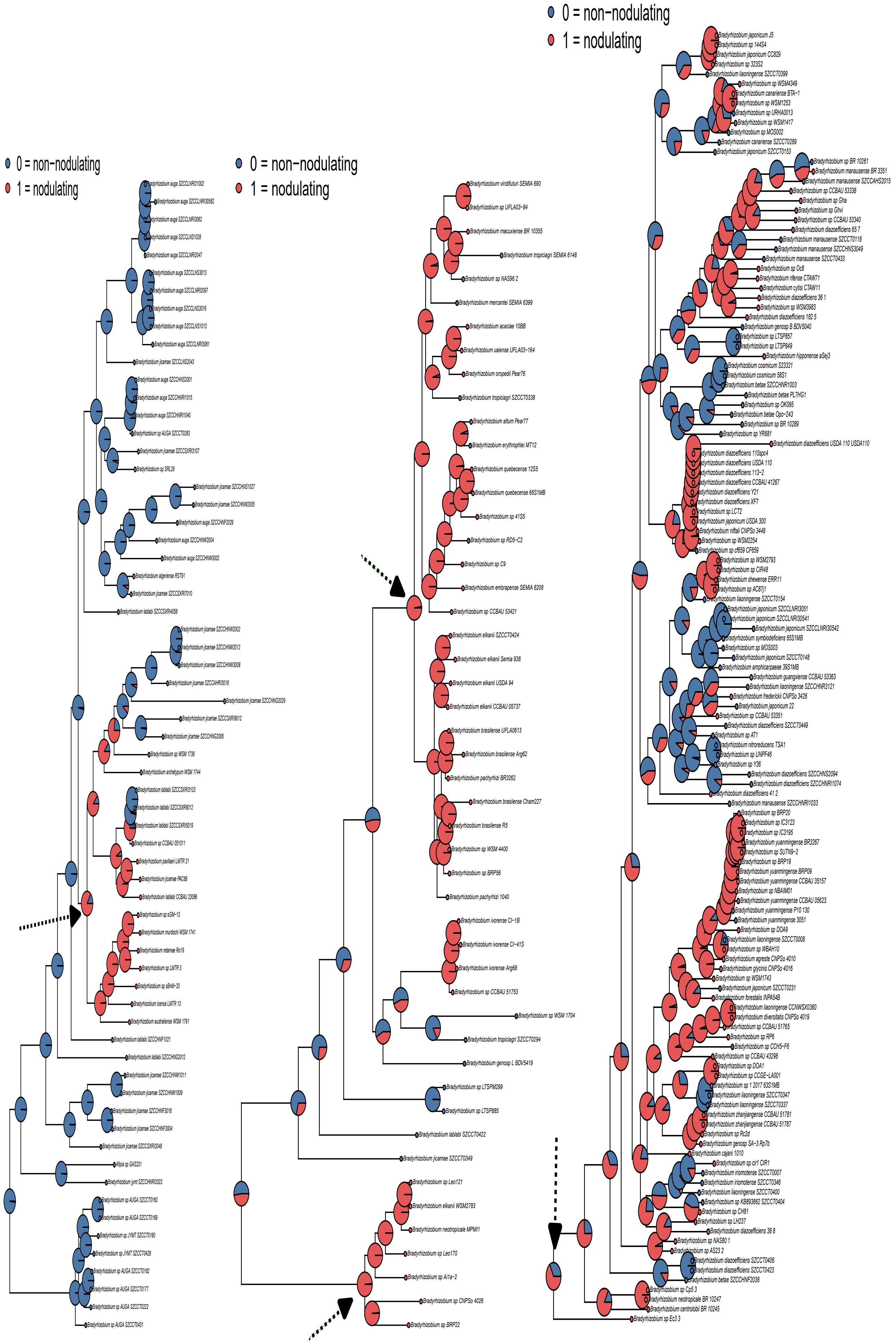

(a) *B. jicamae* supergroup.

(b) *B. elkanii* supergroup.

(c) *B. japonicum* supergroup.

Figure S4: Ancestral state reconstruction of the binary trait, nodulating vs. non-nodulating using the maximum-likelihood method in the R package ape (Paradis and Schliep, 2019). Pie chart shows the marginal posterior probability of the two states at each node. The arrow indicates inferred origins of nodulation.

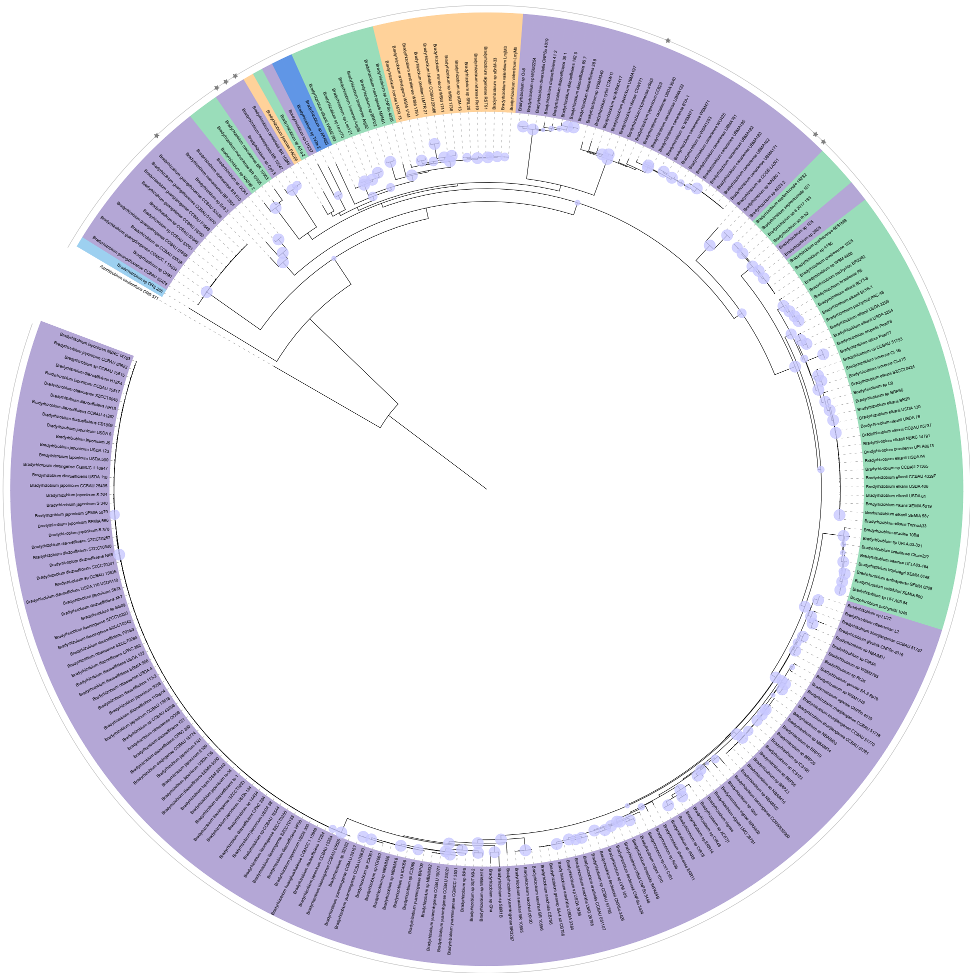

(a) *nodABC* (best-fitting site-homogeneous models)

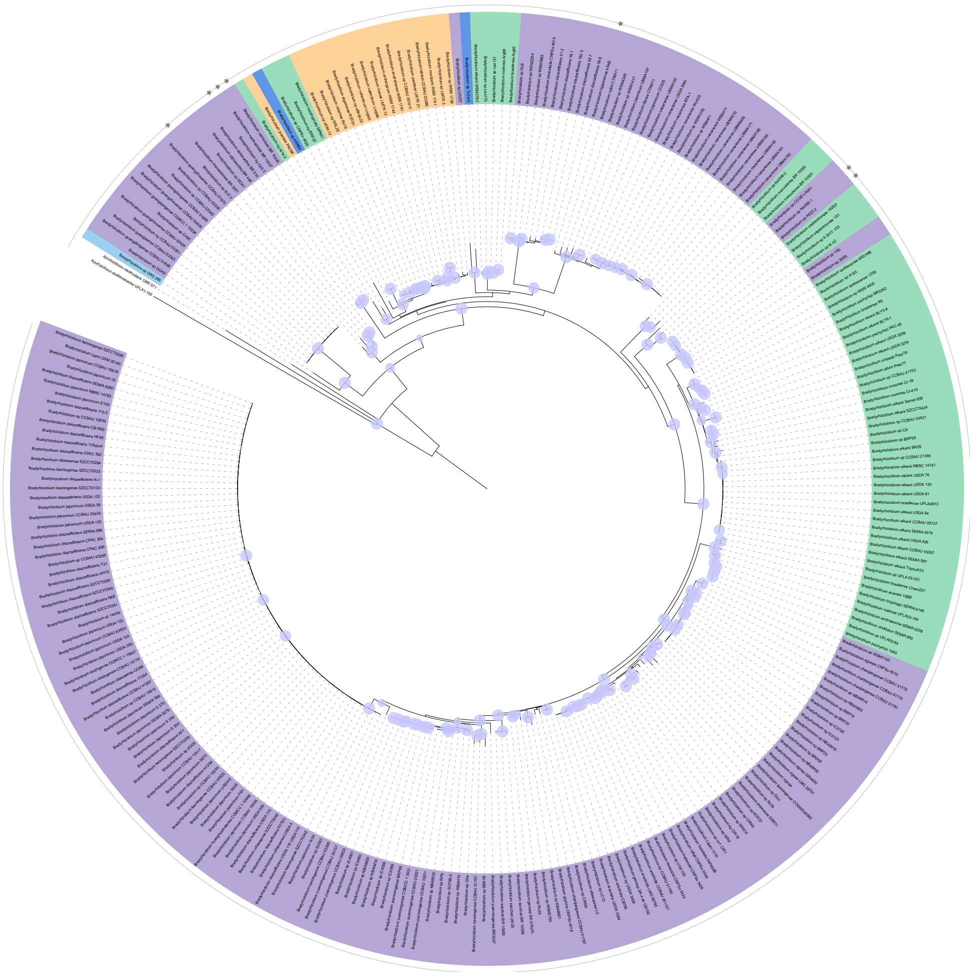

(b) *nodABCIJ* (best-fitting site-homogeneous models)

Figure S5: Phylogenetic trees of *nod* genes under the best-fitting **site-homogeneous substitution models** identified by ModelFinder (Kalyaanamoorthy et al., 2017). Trees are rooted with outgroup sequences of *Azorhizobium*. Colors indicate different *Bradyrhizobium* supergroups, corresponding to those used in Fig. 1. Grey stars in the outer layer indicate basal lineages of nodulating *B. japonicum*. Cyan circles on nodes indicate ultrafast bootstrap values of 90–100, with circle size increasing from smallest to largest.

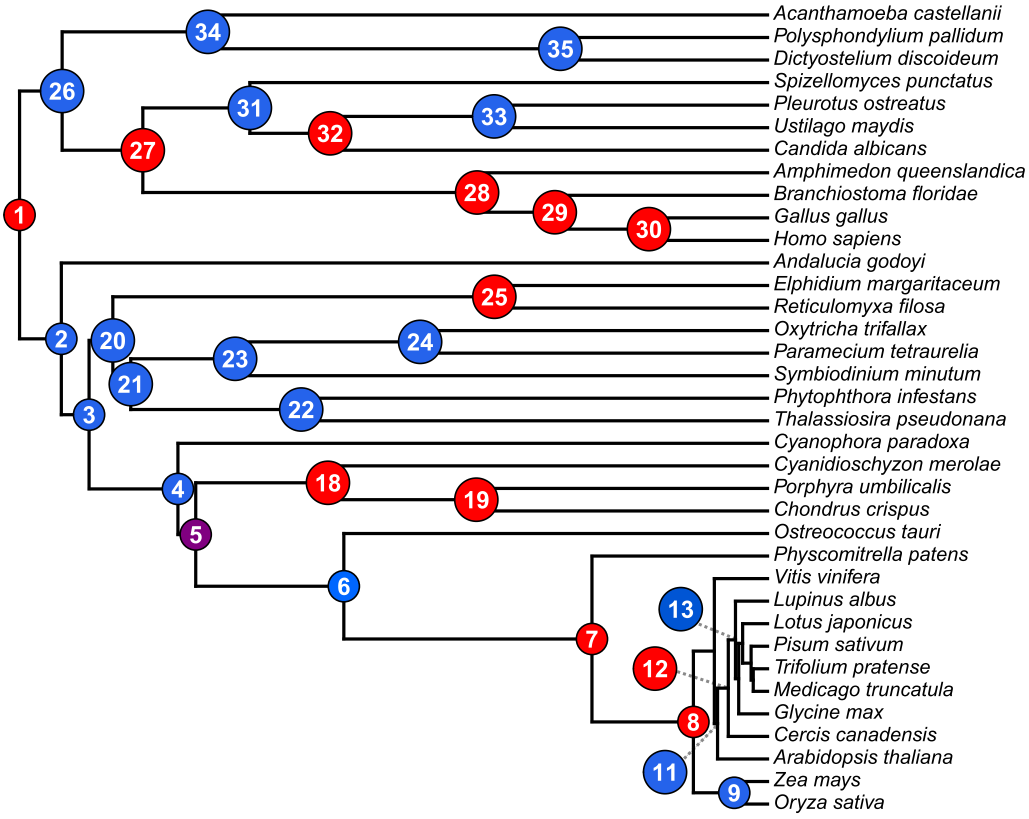

Figure S6: The calibrations used in the eukaryotic mitochondrial clade in molecular dating. Nodes marked with a red square indicate calibrated nodes. The node with a purple circle (node 5) indicates total-group red algae, which replaces node 18 (crown-group red algae) in an alternative analysis reflecting a **more careful evaluation of the fossil records** (see fig. S13). Different calibration settings are detailed in Data S3 and section S2.

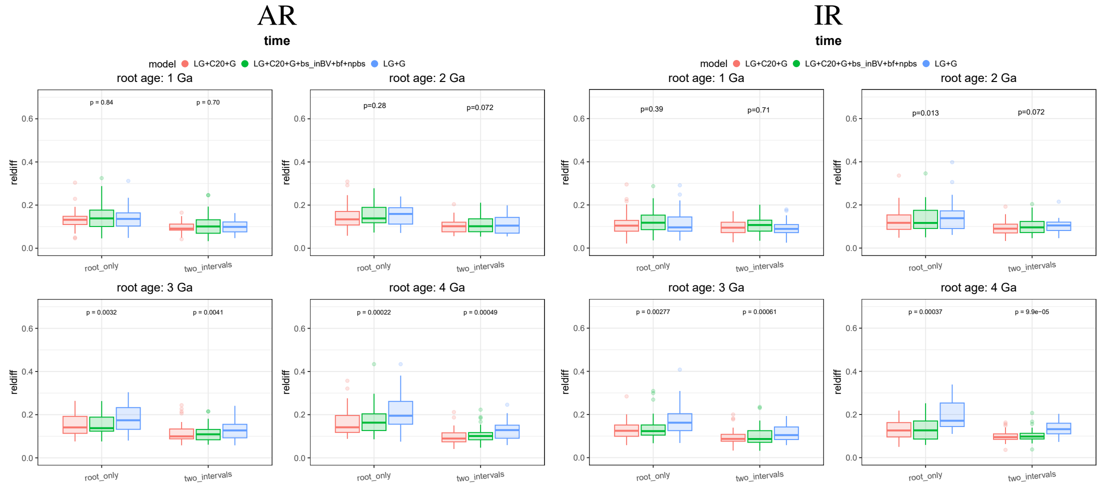

(a) $\lambda = 0.4, \mu = 0.2, \rho = 1$

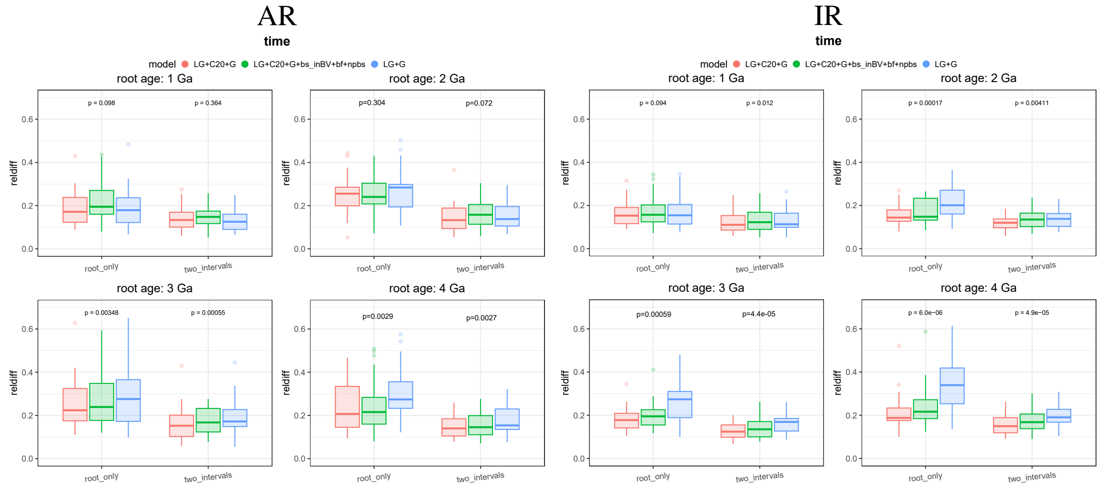

(b) $\lambda = 0.5, \mu = 0.3, \rho = 0.6$

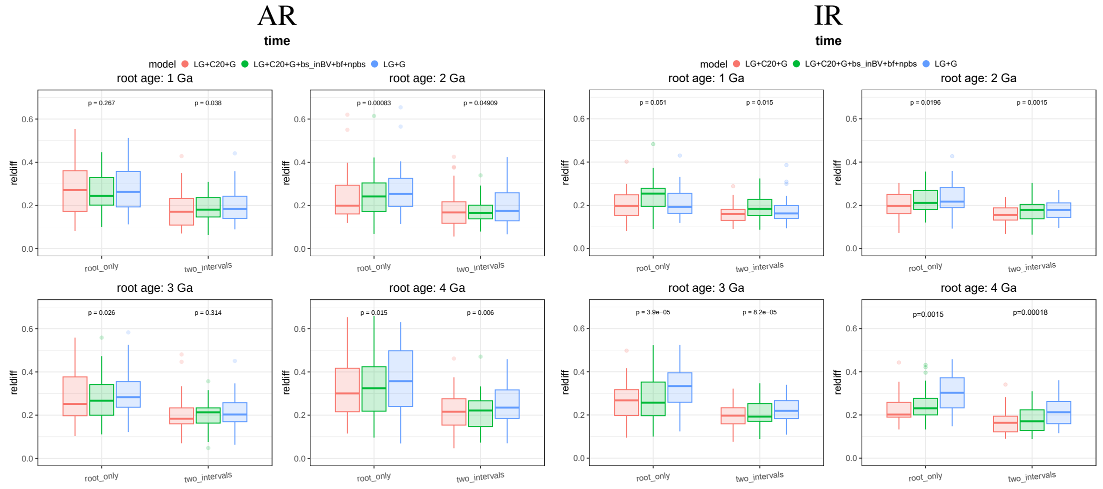

(c) $\lambda = 1.0, \mu = 0.8, \rho = 0.1$

Figure S7: Comparison of **time** estimates by simulated data. Alignments were simulated under LG+C20+G. Time-trees were generated using three birth–death parameter settings mimicking recently branching phylogenies (see also section S1.2). Both autocorrelated-rates (AR) and independent-rates (IR) models were tested. The $y$-axis shows the relative difference (reldiff) between posterior mean node ages and true ages; lower values indicate greater accuracy. Calibration strategies ($x$-axis): ($i$) a single calibration on the root (*root_only*), and ($ii$) one root calibration plus 2 calibrations at internal nodes at the 1/3 and 2/3 age quantiles (*two_intervals*). In all cases, calibration bounds were set to within ± 20% of the true age. LG+C20+G: Hessian estimated by phyloHessian under LG+C20+G; LG+C20+G+bs_inBV+bf+npbs: Hessian estimated by the non-parametric bootstrap (NPBS) approach `bs_inBV` with the nuisance parameter fixed at their best-fitting values in bootstrap; LG+G: Hessian estimated by phyloHessian under LG+G. $P$-values were calculated using paired Wilcoxon signed-rank tests comparing *LG+C20+G* and *LG+G*, followed by Holm adjustment (see section S1.3). Note that although our simulations used absolute root ages ranging from 4.0 to 1.0 Ga, these values should be better interpreted as representing differences in sequence divergence rather than absolute time.

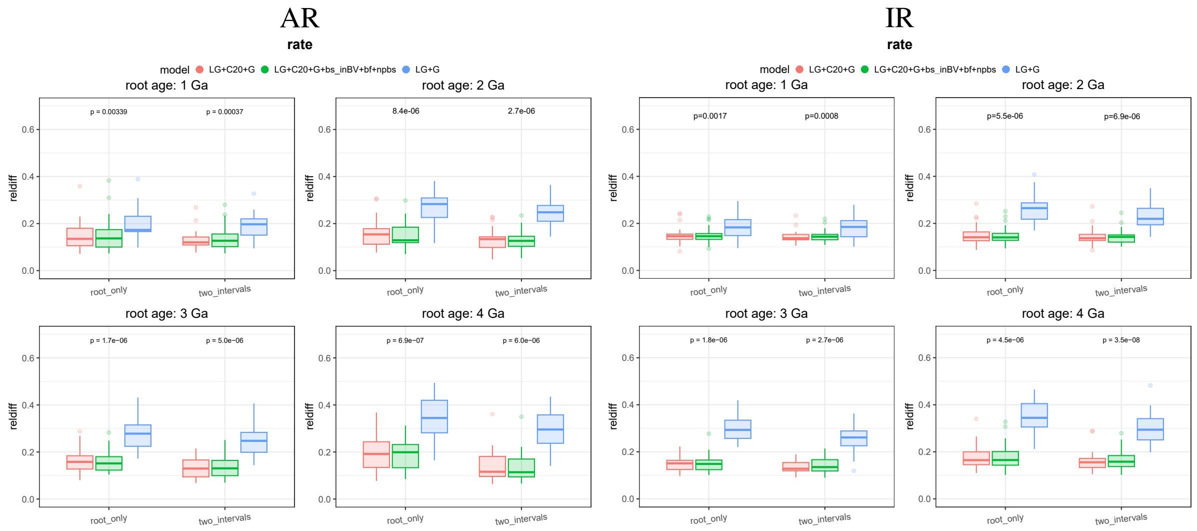

(a) $\lambda = 0.4, \mu = 0.2, \rho = 1$

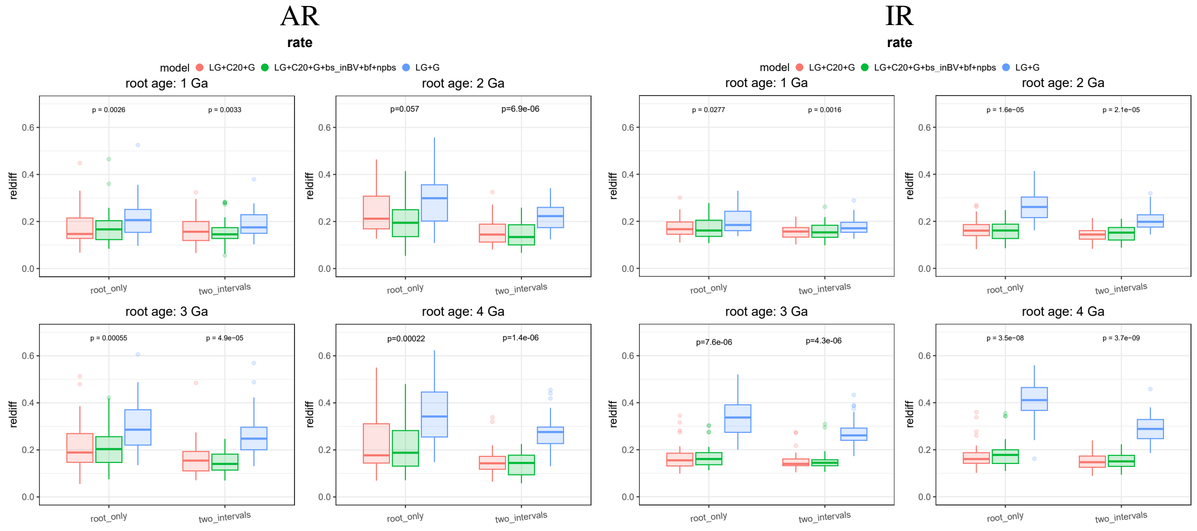

(b) $\lambda = 0.5, \mu = 0.3, \rho = 0.6$

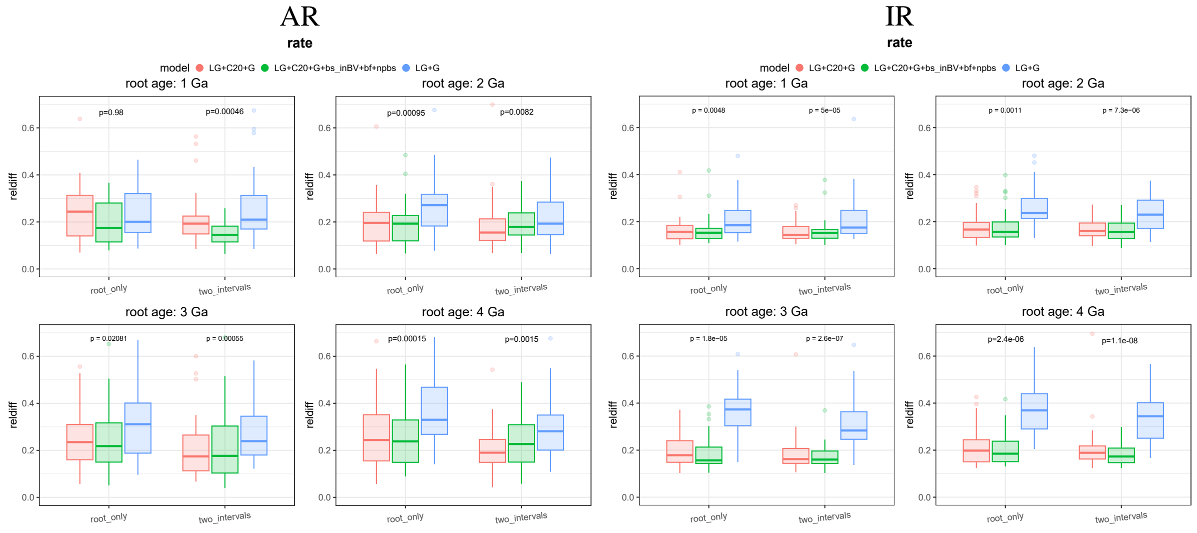

(c) $\lambda = 1.0, \mu = 0.1, \rho = 0.8$

Figure S8: Comparison of **rate** estimates using simulated data with MCMCtree's approximate likelihood method. The caption follows that of fig. S7.

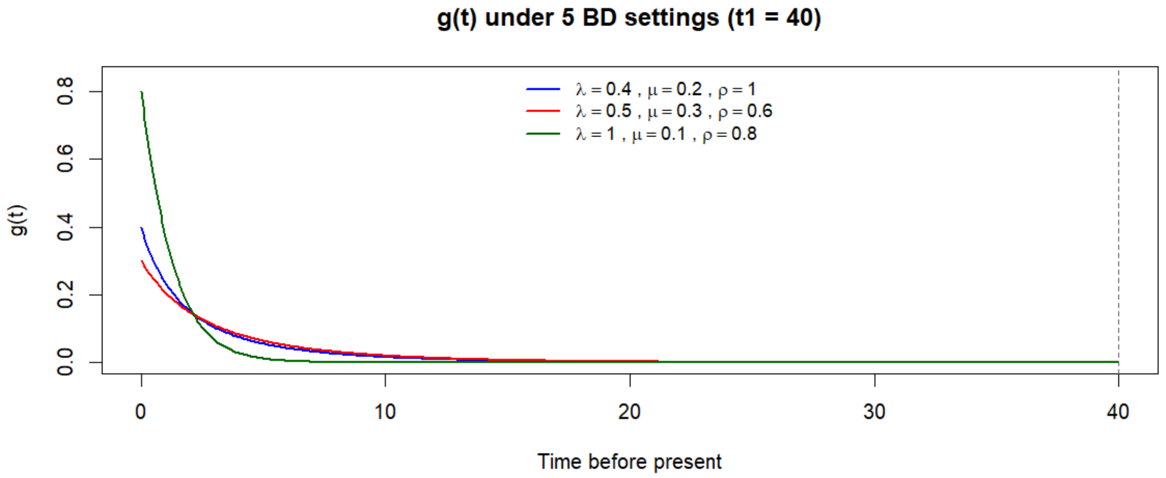

Figure S9: **Density** function $g(t)$ of node ages under a birth–death tree-generating process for the three parameter combinations that **represent recently diverged phylogenies**. The birth–death process is parameterized by the per-lineage birth rate $\lambda$, per-lineage death rate $\mu$, and sampling fraction $\rho$. Differences among curves reflect how alternative birth–death process parameters place relatively a higher probability density on older or younger internal nodes. Blue, red, and green curves reflect recently-diverged phylogenies and denote the settings used in the present study (figs. S7 and S8) where branching density increases toward the present. The $x$-axis shows time before present units: 100 Ma; root age = 4 Ga, and the $y$-axis shows the corresponding value of $g(t)$. Additional details are provided in **section S1.2.1.**

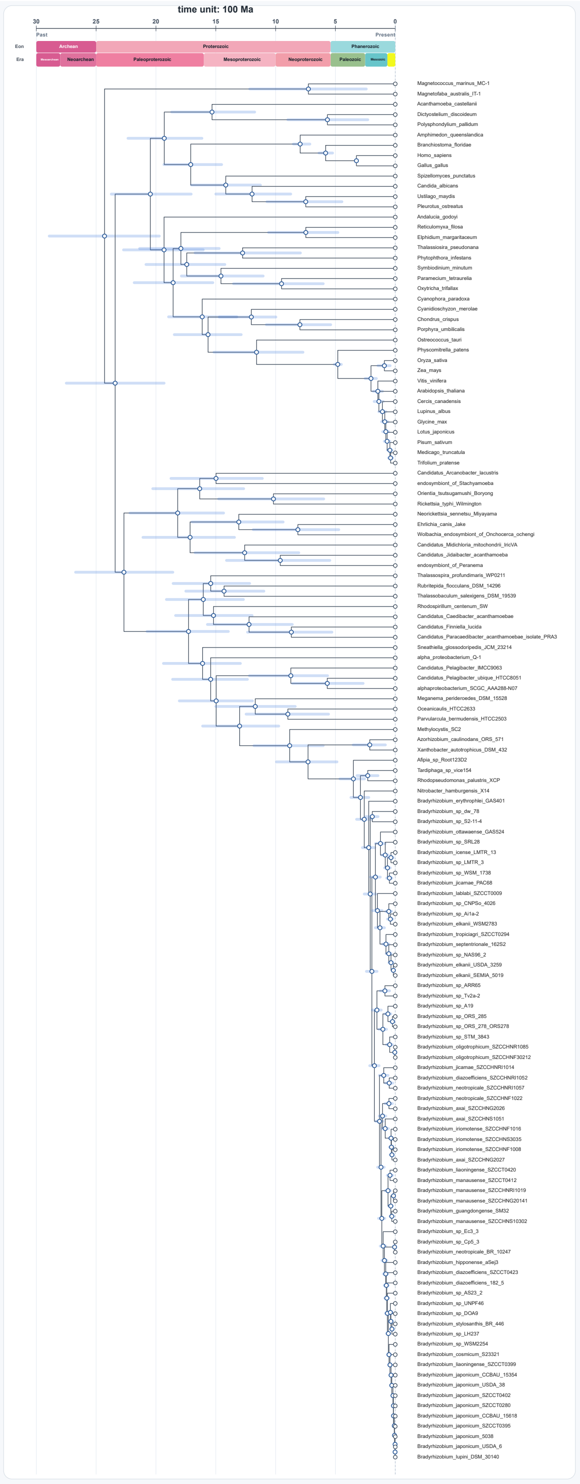

(a) LG+C60+G (PMSF, single partition)

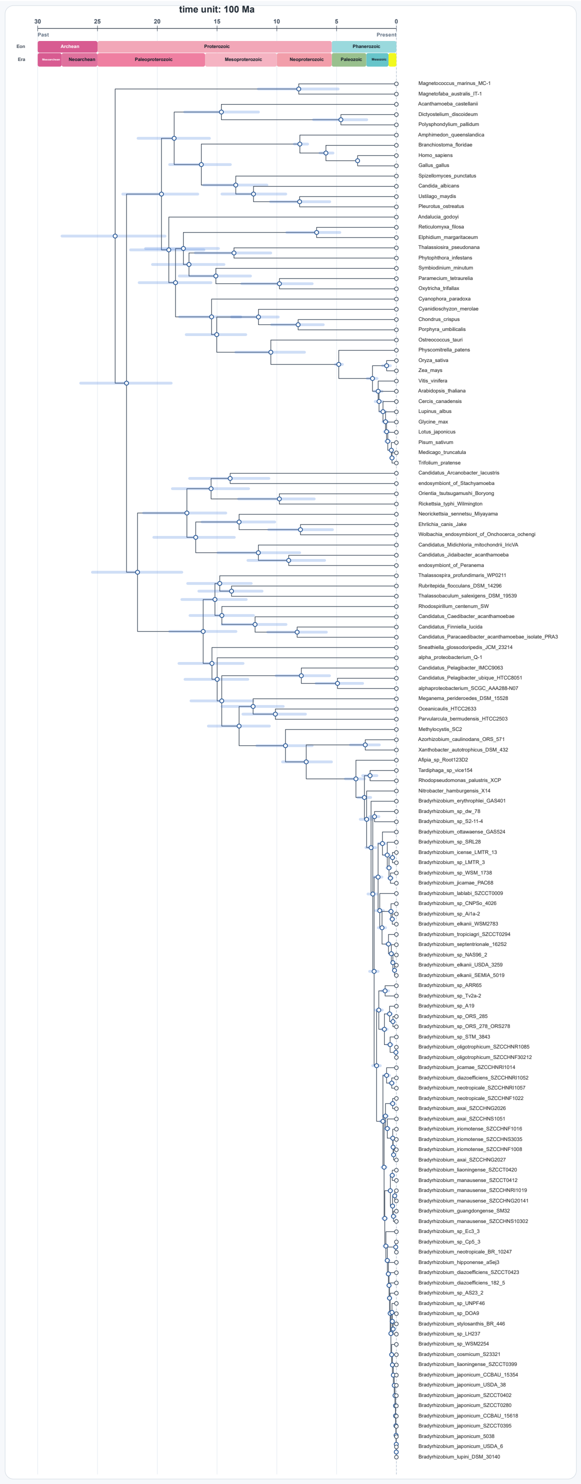

(b) LG+C60+G (PMSF, two partitions)

Figure S10: Chronograms of *Bradyrhizobium* calibrated by the mitochondrial endosymbiosis-based strategy with tip names. Fossil information is given in fig. S6 and Data S3. The chronogram is displayed using `chronogram_viewer`.

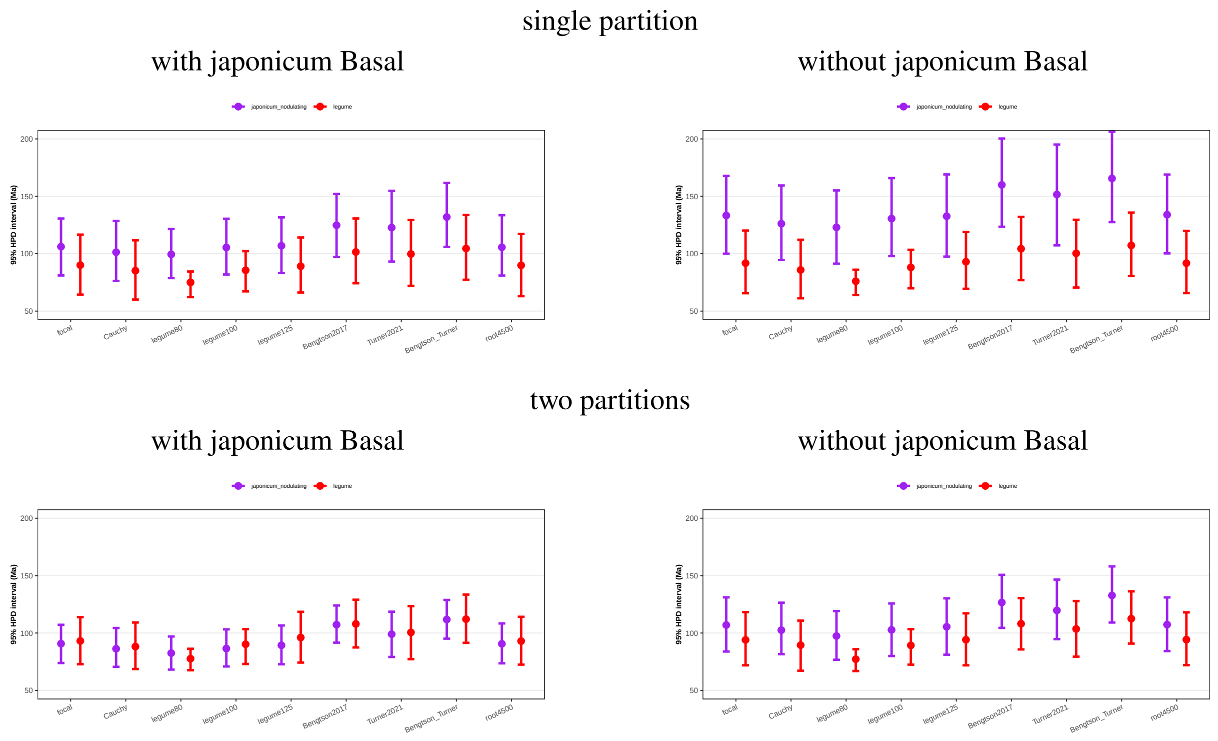

(a) **LG+C20+G (PMSF)**

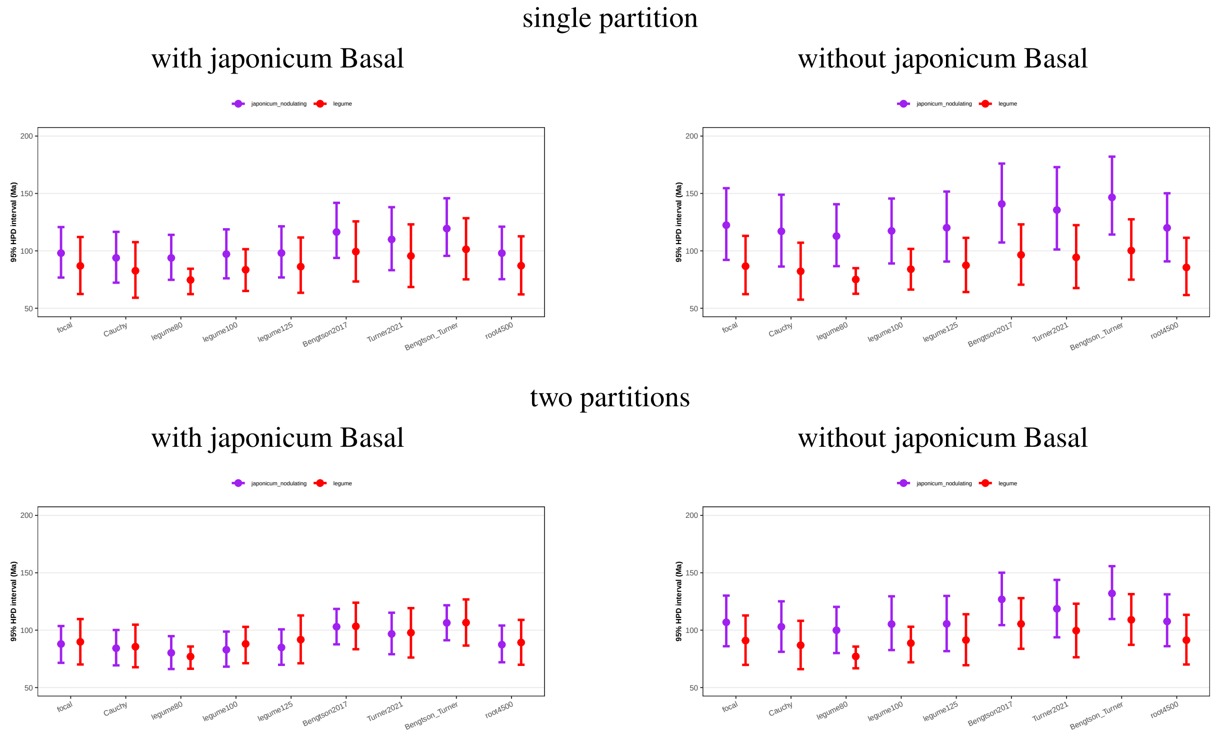

(b) **LG+C60+R+I (PMSF)**

Figure S11: Comparison across calibration schemes of the 95% HPD interval for the posterior divergence time of the crown-group nodulating *B. japonicum* and nodulating legumes under various substitution models. Basal non-nodulating *B. japonicum* lineages are either included (left) or excluded (right), alongside the corresponding estimates for nodulating legumes. The center corresponds to the posterior mean divergence-time estimate. MCMCtree is run using either a single alignment as one partition or the same alignment split into two partitions. a) *LG+C20+G*: substitution model LG with Gamma across-site rate heterogeneity and 20 profile mixtures C20. b) *LG+C60+R+I*: similar to LG+C60+G but with distribution-free (Soubrier et al., 2012; Yang, 1995), instead of Gamma distributed, across-site rate heterogeneity. See also Data S3, Note S2 and Fig. 2 for a detailed description of different calibration schemes.

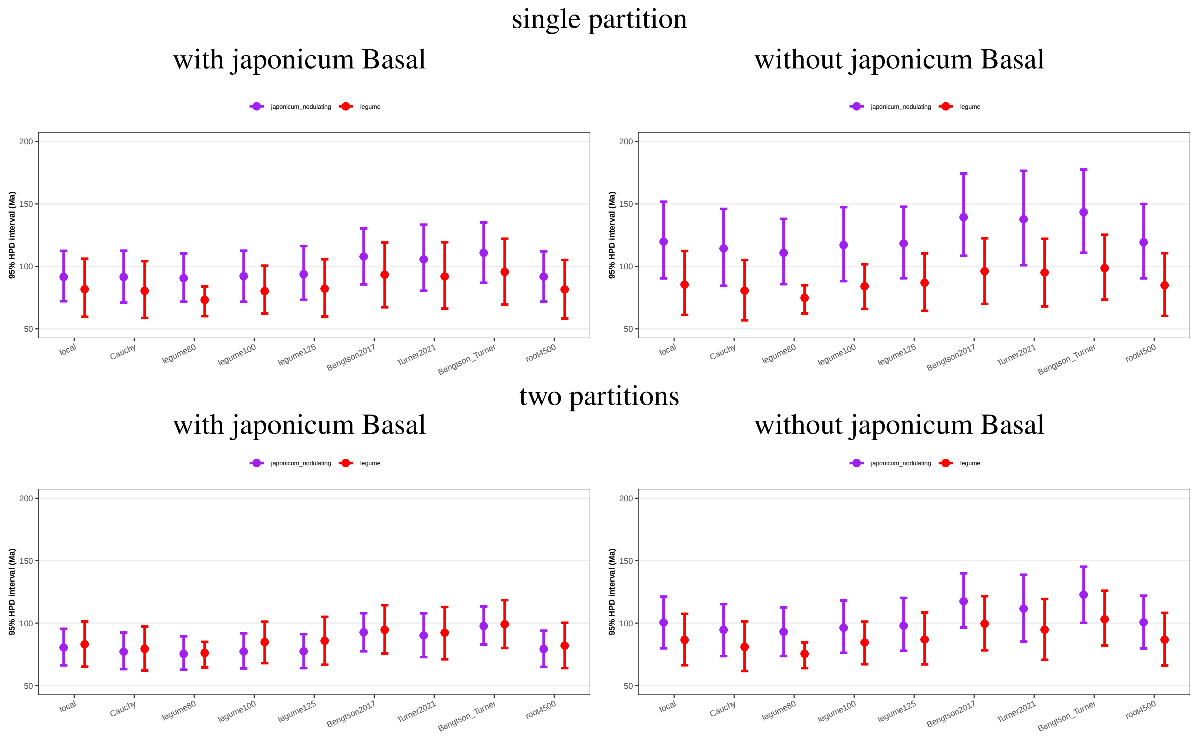

(a) LG+C60+G (PMSF)

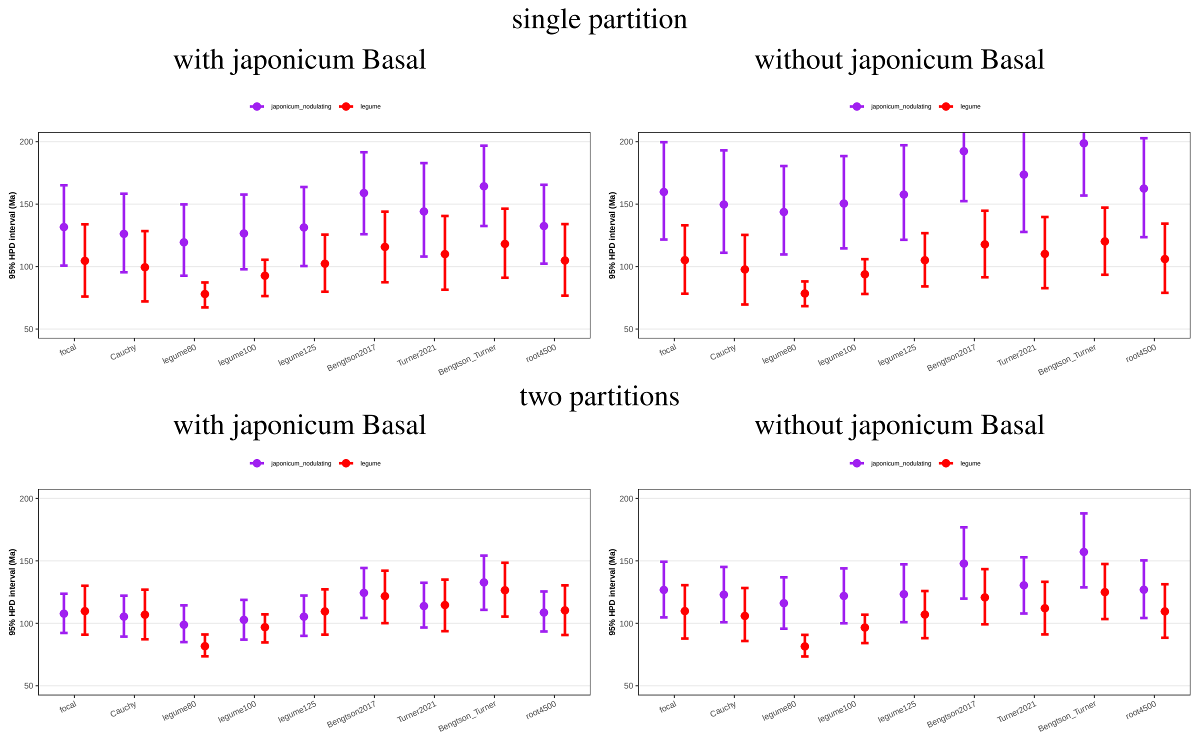

(b) LG+G

Figure S12: Comparison of the 95% HPD intervals for posterior divergence times across calibration schemes of the crown group of nodulating *B. japonicum* and nodulating legumes under different substitution models using the full gene set *gene53* (see table S1).

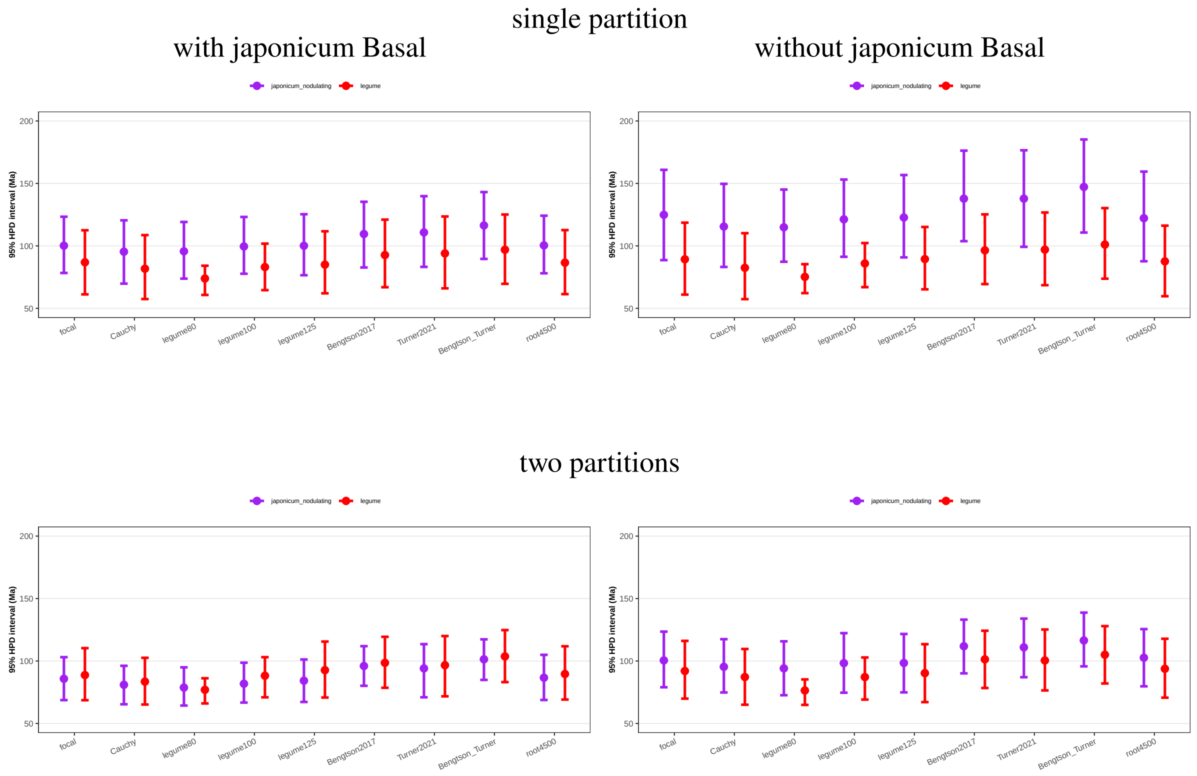

Figure S13: Comparison across calibration schemes of the 95% HPD interval for the posterior divergence time of the crown group of nodulating *B. japonicum* and legumes under the site-heterogeneous mixture substitution model LG+C60+G (PMSF) with a more conservative phylogenetic placement of the red fossil records. Different from figs. S11 and S12 as well as Fig. 2, the calibration on the crown-group red algae (Node 18 in fig. S6) is placed to its **total group** (Node 5 in fig. S6). The replacement of the calibration reflects a more careful and conservative evaluation of the phylogenetic position of fossils (section S2.2.7).

### Supplementary Notes

#### S1 Additional notes on molecular clock dating

##### S1.1 Improved bootstrap-based approach for using mixture substitution models in MCMCtree

MCMCtree uses an approximate likelihood that summarizes the log-likelihood surface around the branch-length maximum-likelihood estimates (MLEs). Let $\boldsymbol{\theta}$ denote the vector of branch lengths and $\hat{\boldsymbol{\theta}}$ its MLE under a fixed topology and substitution model. Following dos Reis and Yang (2011), the log-likelihood is approximated near $\hat{\boldsymbol{\theta}}$ by

$$\ell(\boldsymbol{\theta}) \approx \ell(\hat{\boldsymbol{\theta}}) + \boldsymbol{g}^\top(\boldsymbol{\theta} - \hat{\boldsymbol{\theta}}) + \frac{1}{2}(\boldsymbol{\theta} - \hat{\boldsymbol{\theta}})^\mathsf{T}\mathbf{H}(\hat{\boldsymbol{\theta}})(\boldsymbol{\theta} - \hat{\boldsymbol{\theta}}), \tag{1}$$

where $\boldsymbol{g}$ and $\mathbf{H}(\hat{\boldsymbol{\theta}})$ are the gradient and Hessian with respect to branch lengths. At the MLE, the gradient is zero except for zero-length branches (dos Reis and Yang, 2011), so the approximate likelihood is determined mainly by $\hat{\boldsymbol{\theta}}$ and $\mathbf{H}(\hat{\boldsymbol{\theta}})$.

In PAML, the Hessian is computed numerically under substitution models implemented in PAML (dos Reis and Yang, 2011; Thorne et al., 1998). This limits MCMCtree dating to substitution models available in PAML.

Our bootstrap-based approach bypasses this limitation by estimating the likelihood curvature from bootstrap variation in branch-length estimates, instead of the finite-difference numerical method. Specifically, following Wang and Luo (2025), we approximate

$$\mathbf{H}(\hat{\boldsymbol{\theta}}) \approx -\mathbf{V}(\hat{\boldsymbol{\theta}})^{-1}, \tag{2}$$

where $\mathbf{V}(\hat{\boldsymbol{\theta}})$ is the covariance matrix of the branch-length estimator. Since this covariance matrix can be estimated from bootstrap branch lengths inferred under any substitution model, the resulting approximate likelihood can be used in MCMCtree without requiring that model to be implemented in CODEML or any other similar software (Demotte et al., 2025; Wang and Meade, 2026). Note however that this holds only if the model is correctly specified. Under model misspecification the sandwich estimator should be used (White, 1982) and the case is more complex and beyond the scope of the present study.

###### S1.1.1 Fixing nuisance parameters at their best-fitting values

Here, we applied the following improvements to the bootstrap-based method `bs_inBV` developed in Wang and Luo (2025).

To better match MCMCtree's approximate-likelihood implementation, nuisance parameters, such as $\alpha$ in +G or the weights of Cxx mixture components, are now fixed at their MLEs during MCMC updates. Let the full parameter vector be $\boldsymbol{\Theta} = (\boldsymbol{\theta}, \boldsymbol{\eta})$, where $\boldsymbol{\theta}$ denotes branch lengths and $\boldsymbol{\eta}$ denotes nuisance parameters such as the rate-heterogeneity parameters ($\alpha$ in the +G model) and the weights of mixture components in a mixture model. If $(\hat{\boldsymbol{\theta}}, \hat{\boldsymbol{\eta}})$ are the MLEs from the original alignment, then bootstrap replicates were analyzed conditional on $\hat{\boldsymbol{\eta}}$, yielding branch-length estimates $\hat{\boldsymbol{\theta}}^{(b)}$.

The covariance matrix was estimated as

$$\widehat{\mathbf{V}}_{\text{fix}}(\hat{\boldsymbol{\theta}} \mid \hat{\boldsymbol{\eta}}) = \frac{1}{B-1}\sum_{b=1}^{B}\left(\hat{\boldsymbol{\theta}}^{(b)} - \bar{\hat{\boldsymbol{\theta}}}\right)\left(\hat{\boldsymbol{\theta}}^{(b)} - \bar{\hat{\boldsymbol{\theta}}}\right)^\mathsf{T}, \tag{3}$$

where

$$\bar{\hat{\boldsymbol{\theta}}} = \frac{1}{B}\sum_{b=1}^{B}\hat{\boldsymbol{\theta}}^{(b)}. \tag{4}$$

The corresponding Hessian approximation is

$$\mathbf{H}_{\text{fix}}(\hat{\boldsymbol{\theta}} \mid \hat{\boldsymbol{\eta}}) \approx -\widehat{\mathbf{V}}_{\text{fix}}(\hat{\boldsymbol{\theta}} \mid \hat{\boldsymbol{\eta}})^{-1}. \tag{5}$$

In the older versions of `bs_inBV` (Wang and Luo, 2025), $\mathbf{H}(\hat{\boldsymbol{\theta}})$ was approximated as the negative inverse of the $\boldsymbol{\theta}$-submatrix of $\mathbf{V}(\boldsymbol{\Theta})$. This submatrix is generally not the same as $\mathbf{V}_{\text{fix}}(\hat{\boldsymbol{\theta}} \mid \hat{\boldsymbol{\eta}})$. Alternatively, the required Hessian matrix, i.e. $\mathbf{H}(\hat{\boldsymbol{\theta}})$, can be calculated by Schur's complement

$$\mathbf{H}(\hat{\boldsymbol{\theta}}) \approx -\left(\mathbf{V}_{\theta\theta} - \mathbf{V}_{\theta\eta}\mathbf{V}_{\eta\eta}^{-1}\mathbf{V}_{\eta\theta}\right)^{-1}, \tag{6}$$

which does not need to fix the nuisance parameters at their MLEs.

##### S1.1.2 Regularization of singular covariance matrices

For some datasets, $\widehat{\mathbf{V}}_{\text{fix}}(\hat{\boldsymbol{\theta}} \mid \hat{\boldsymbol{\eta}})$ was singular, thus non-invertible. This usually happens when one or more branch lengths were estimated as zero or at the minimum value across all bootstrap alignments (Wang and Luo, 2025), particularly common for super-short branches (e.g., in the presence of completely identical sequences). We regularized it via eigendecomposition:

$$\widehat{\mathbf{V}}_{\text{fix}} = \mathbf{Q}\boldsymbol{\Lambda}\mathbf{Q}^{\mathsf{T}}, \qquad \widetilde{\boldsymbol{\Lambda}} = \text{diag}(\max(\lambda_1, \varepsilon), \ldots, \max(\lambda_n, \varepsilon)), \tag{7}$$

with $\varepsilon = 10^{-6}$, and then set

$$\mathbf{H}_{\text{fix}}(\hat{\boldsymbol{\theta}} \mid \hat{\boldsymbol{\eta}}) \approx -\left(\mathbf{Q}\widetilde{\boldsymbol{\Lambda}}\mathbf{Q}^{\mathsf{T}}\right)^{-1}. \tag{8}$$

#### S1.2 Evaluating the performance of different substitution models for recently diverged phylogenies

##### S1.2.1 Choice of birth–death process parameters

To mimic the diversification pattern of the *Bradyrhizobium* dataset, in which most lineages are young relative to the root, we selected birth–death time-prior parameters that favor recent branching. Following Yang and Rannala (2006) (Eqs. 4–10), the density of internal node ages ($\lambda \neq \mu$) is described by

$$g(t) = \frac{\lambda p_1(t)}{v_{t_1}},$$

where

$$p_1(t) = \frac{1}{\rho}P(0,t)^2 e^{(\mu-\lambda)t}, \qquad v_{t_1} = 1 - \rho^{-1}P(0,t_1)e^{(\mu-\lambda)t_1},$$

and

$$P(0,t) = \frac{\rho(\lambda-\mu)}{\rho\lambda + [\lambda(1-\rho)-\mu]e^{(\mu-\lambda)t}}.$$

This kernel defines the probability distribution of internal node ages (time prior) conditional on the root age $t_1$. The selected parameter sets favor phylogenies with node ages concentrated toward the present, consistent with shallow or recently diverged trees. Three sets of parameters were chosen: $\lambda = 0.4, \mu = 0.2, \rho = 1$, $\lambda = 0.5, \mu = 0.3, \rho = 0.6$ and $\lambda = 1.0, \mu = 0.1, \rho = 0.8$. Their probability density plot of the branching is given in fig. S9.

##### S1.2.2 Simulation procedure

True root ages were set to 1.0, 2.0, 3.0, and 4.0 Ga, following Wang and Luo (2025); Wang and Meade (2026). For each root age, we simulated 30 birth–death timetrees with 20 tips using `TreeSim` (Stadler, 2011), with parameters given in section S1.2.1. Branch-specific rates were then simulated under independent-rates (IR) and autocorrelated-rates (AR) models. For IR, rates followed a lognormal distribution with $\mu_{\text{IR}} = -3.7$ and $\sigma_{\text{IR}} = 0.2$, corresponding to a mean of approximately 0.025 substitutions per site per 100 Myr and a standard deviation of approximately 0.005 (Wang and Meade, 2026). AR parameters were chosen following Wang and Meade (2026) so that the expected sample mean and variance of log-rates matched those under the IR model.

For each simulated timetree, we generated a 300-site amino acid alignment using `AliSim` (Ly-Trong et al., 2022) under LG+C20+G4{1.0}, combining the LG exchangeability matrix, the C20 profile mixture model (Le et al., 2008), and discrete-gamma among-site rate variation with four categories and $\alpha = 1.0$.

#### S1.3 Comparison between substitution models and approaches to estimating the Hessian

##### S1.3.1 Relative difference metric

Following Wang and Meade (2026), we assessed the performance of different substitution models using the relative difference (*reldiff*), which quantifies how closely posterior mean estimates from `MCMCtree` matched the true values used in simulation. For a given model $m$, *reldiff* was defined as

$$\text{reldiff} = \frac{1}{n} \sum_{i=1}^{n} \frac{|b_i^m - b_i^{\text{true}}|}{\max(b_i^m,\, b_i^{\text{true}})}, \tag{9}$$

where $b_i^m$ denotes the posterior mean divergence time or branch-specific substitution rate inferred under model $m$ for node or branch $i$, $b_i^{\text{true}}$ is the corresponding true simulated value, and $n$ is the number of divergence times or branch rates being compared. Lower values indicate a more accurate inference.

##### S1.3.2 Simulation settings and Hessian-estimation methods

All alignments were simulated under the mixture model LG+C20+G4{1.0} (discrete Gamma distribution with $\alpha = 1$). We applied two substitution models in calculating the gradient and Hessian matrix, LG+C20+G (right model) and LG+G (wrong model). We considered the following different Hessian-estimation methods for both models:

- *LG+G, LG+C20+G*: outer product of scores (OPS) method, based on the outer product of the first-order derivative of the log-likelihood (default in phyloHessian and CODEML v4.3+);
- *+bs_inBV+bf+npbs*: Hessian estimated by the approach based on non-parametric bootstrap (NPBS) with nuisance parameters fixed at best-fitting (bf) values (i.e., MLEs) during bootstrap (section S1.1).

##### S1.3.3 Effects of substitution-model choice and Hessian approximation

As shown in fig. S7, when the true root age was over 3.0-2.0 Ga, the mixture model LG+C20+G consistently outperformed LG+G. Branch-specific posterior rates were also more accurately estimated under LG+C20+G (fig. S8) in divergence time estimates. The pattern was more obvious for the rate estimates, likely because calibrations constrain node times more strongly than rates (Wang and Meade, 2026). Our results reveal a general pattern: the impact of substitution-model choice on molecular dating estimates increases with sequence divergence. This pattern agrees with previous studies (Wang and Luo, 2025; Wang and Meade, 2026) and highlights the importance of appropriate substitution models for deep-time dating. The bootstrap-based approaches in `bs_inBV` performed comparably to the finite-difference OPS method figs. S7 and S8. This suggests that the bootstrap-based method is a useful alternative to finite-difference Hessian calculation (dos Reis and Yang, 2011), particularly for substitution models for which direct Hessian computation is not available.

### S2 Time-Calibrations

#### S2.1 Focal calibrations

For the focal calibration strategy, we basically followed Wang and Luo (2021) for the calibrations in eukaryotes for a combined dataset of the mito- and nuclear-encoded gene sets in that study. For simplicity, we suggest readers reference the original study for these calibrations used in the focal calibration scheme.

We additionally included a fossil-based calibration for the **total group** of nodulating legumes (i.e., the crown group of Caesalpinioideae and Papilionoideae) on Node 12 (fig. S6). We constrained its minimum age to be 60 Ma based on the fossils which are believed to belong to the subfamily Caesalpinioideae (Herendeen and Crane, 1992; Herrera et al., 2019). In that study, they are compression/impression fossils of legume fruits and leaves from the Middle–Late Paleocene of Colombia. These fossils have also been used in other molecular dating studies (Lavin et al., 2005; Magallon and Sanderson, 2001).

#### S2.2 Alternative calibrations

##### S2.2.1 *Bengtson2017*

In the alternative calibration scheme *Bengtson2017*, we constrained the minimum age of crown-group red algae to 1.6 Ga, a relatively aggressive calibration. This was based on the fossils *Ramathallus lobatus* and *Rafatazmia chitrakootia* from the Vindhyan Supergroup, which were proposed to represent crown red algae (Bengtson et al., 2017). However, because this interpretation was questioned by several other studies (Betts et al., 2018; Gibson et al., 2018) and was believed to be highly controversial, we considered this calibration only as an alternative scenario to assess its effect on divergence-time estimates and the relative divergence order of *Bradyrhizobium* and legumes.

##### S2.2.2 *Turner2021*

In the alternative calibration scheme *Turner2021*, we constrained the minimum age of crown-group animals to 890 Ma, based on putative sponge fossils from deep Proterozoic rocks (Turner, 2021). This interpretation relied on vermiform structures being identified as spongin fibres of keratosan sponges, but it has been disputed by later studies (Carlisle et al., 2024; Kris and McMenamin, 2021; Rossi et al., 2026), which suggested alternative explanations and called for further evidence. Because this interpretation remains controversial, we considered this calibration only as an alternative scenario to assess its effect on molecular dating.

###### S2.2.3 *Bengtson_Turner*

The combination of the alternative settings of *Bengtson2017* and *Turner2021*.

###### S2.2.4 *root4500*

We alternatively set the root maximum to 4500 Ma instead of the 3000 Ma maximum used in the focal scheme, roughly the age of Earth. This was to test if the impact of an arbitrarily set root maximum could impact the divergence time estimate. As expected, the results remained highly similar (Fig. 3), because of the maximum age constraint imposed at many internal nodes, consistent with our previous works (Liao et al., 2024; Wang and Luo, 2021, 2025).

###### S2.2.5 *Cauchy*

For calibrations lacking a closely constrained maximum age, namely those whose maximum bound had been set by the 1.891-Ga fossil, we removed the hard maximum bound and instead applied a truncated Cauchy distribution in `MCMCTree` (see Data S3). Specifically, the prior was specified as $T \sim L(t_L, p, c, p_L)$, where $t_L$ is the minimum fossil age, $p$ determines how far the mode lies above the minimum bound, $c$ controls how sharply the distribution decays, and $p_L$ is the left-tail probability that the minimum bound is violated. Here, we used the default parameters in `MCMCtree` $p = 0.1$, $c = 0.1$, and $p_L = 0.025$.

###### S2.2.6 *legume80, legume100, legume125*

We also explored alternative maximum bounds to the total group of nodulating legumes (i.e., LCA of *Glycine max* and *Lupinus*; fig. S6, Node 13). This is based on secondary calibrations from previous studies for *legume80* (Hohmann et al., 2015) and *legume100* (Foster et al., 2017) which estimated the node's age to be 80 Ma and 100 Ma respectively, and the minimum age bound for the total-group dicots for *legume125* (125 Ma; see also Data S3).

###### S2.2.7 Total vs. crown group of red algae

Total-group red algae: While in the main analysis we placed the calibration to the crown group of red algae (fig. S6, Node 18), a more conservative phylogenetic placement of the corresponding fossil records is probably the total group. We alternatively constrained the total group, instead of the crown group, of red algae's minimum age to be $1.033$ Ga according to the same fossil records (Turner and Kamber, 2012). The time estimates resulting from using this alternative calibration scheme are displayed in fig. S13.
